# Rational Inattention and Tonic Dopamine

**DOI:** 10.1101/2020.10.04.325175

**Authors:** John G. Mikhael, Lucy Lai, Samuel J. Gershman

## Abstract

Slow-timescale (tonic) changes in dopamine (DA) contribute to a wide variety of processes in reinforcement learning, interval timing, and other domains. Furthermore, changes in tonic DA exert distinct effects depending on when they occur (e.g., during learning vs. performance) and what task the subject is performing (e.g., operant vs. classical conditioning). Two influential theories of tonic DA—the average reward theory and the Bayesian theory in which DA controls precision—have each been successful at explaining a subset of empirical findings. But how the same DA signal performs two seemingly distinct functions without creating crosstalk is not well understood. Here we reconcile the two theories under the unifying framework of ‘rational inattention,’ which (1) conceptually links average reward and precision, (2) outlines how DA manipulations affect this relationship, and in so doing, (3) captures new empirical phenomena. In brief, rational inattention asserts that agents can increase their precision in a task (and thus improve their performance) by paying a cognitive cost. Crucially, whether this cost is worth paying depends on average reward availability, reported by DA. The monotonic relationship between average reward and precision means that the DA signal contains the information necessary to retrieve the precision. When this information is needed after the task is performed, as presumed by Bayesian inference, acute manipulations of DA will bias behavior in predictable ways. We show how this framework reconciles a remarkably large collection of experimental findings. In reinforcement learning, the rational inattention framework predicts that learning from positive and negative feedback should be enhanced in high and low DA states, respectively, and that DA should tip the exploration-exploitation balance toward exploitation. In interval timing, this framework predicts that DA should increase the speed of the internal clock and decrease the extent of interference by other temporal stimuli during temporal reproduction (the central tendency effect). Finally, rational inattention makes the new predictions that these effects should be critically dependent on the controllability of rewards, that post-reward delays in intertemporal choice tasks should be underestimated, and that average reward manipulations should affect the speed of the clock—thus capturing empirical findings that are unexplained by either theory alone. Our results suggest that a common computational repertoire may underlie the seemingly heterogeneous roles of DA.

**Author Summary:** The roles of tonic dopamine (DA) have been the subject of much speculation, partly due to the variety of processes it has been implicated in. For instance, tonic DA modulates how we learn new information, but also affects how previously learned information is used. DA affects the speed of our internal timing mechanism, but also modulates the degree to which our temporal estimates are influenced by context. DA improves performance in some tasks, but seems only to affect confidence in others. Are there common principles that govern the role of DA across these domains? In this work, we introduce the concept of ‘rational inattention,’ originally borrowed from economics, to the DA literature. We show how the rational inattention account of DA unites two influential theories that are seemingly at odds: the average reward theory and the Bayesian theory of tonic DA. We then show how this framework reconciles the diverse roles of DA, which cannot be addressed by either theory alone.

## Introduction

The functions of dopamine (DA) have been a subject of debate for several decades, due in part to the bewildering variety of processes in which it participates. DA plays diverse roles in reinforcement learning and action selection [1–9], motor control [10–12], vision [13, 14], interval timing [15–18], and attention [19]. Furthermore, DA functions through at least two channels: fast-timescale (phasic) signals and slow-timescale (tonic) signals [20]. While a large body of evidence has shown that phasic DA corresponds remarkably well with a ‘reward prediction error’ in reinforcement learning models [1–3, 21, 22], the functions of tonic DA, which span the domains above, remain unclear. Does this diversity of function reflect a heterogeneous computational repertoire? Or are there common principles that govern the role of tonic DA across these domains?

Across both experiments and theory, tonic DA is most widely studied in the domains of reinforcement learning and decision making. Experiments with Parkinson’s patients and healthy subjects have suggested that, when learning the associations between actions (or stimuli) and rewards, high DA facilitates learning from positive feedback, and low DA facilitates learning from negative feedback across a variety of tasks [4, 5, 23–27]. On the other hand, when this learned information must subsequently be used to select actions, DA seems to control the exploration-exploitation trade-off: Here, high DA promotes exploitation of actions with higher learned value and increases motivation. Low DA, on the other hand, promotes exploration [28–30] (but see [31, 32] and Discussion) and decreases motivation [33–35]. Recently developed computational models allow DA to achieve both learning and performance roles by endowing it with separate computational machinery for each [27, 36]. However, whether and how the effects of DA during learning and performance may be related at an algorithmic level remain open questions.

Another well-studied role of DA relates to its influence on interval timing. Much like in reinforcement learning, DA also seems to have two broad effects here. First, DA seems to modulate the speed of the internal timing mechanism: Acutely increasing DA levels results in behaviors consistent with a faster ‘internal clock,’ and acutely decreasing DA levels results in a slower internal clock [15–17, 37, 38]. In addition, unmedicated Parkinson’s patients, who are chronically DA-depleted, exhibit a distinct timing phenomenon known as the ‘central tendency’: When these patients learn intervals of different durations, they tend to overproduce the shorter intervals and underproduce the longer intervals [39–41]. While the central tendency has been observed in healthy subjects [42–46] and animals [47], it is most pronounced in unmedicated Parkinson’s patients, and DA repletion in these patients rescues veridical timing [39]. How the effects of DA on the speed of the internal clock and the central tendency are related, if at all, remains unclear.

Reinforcement learning and interval timing, though related at a theoretical and neural level [22, 48], have, until recently, largely been treated separately [49–51]. This has led to separate theories of DA developing for each: In reinforcement learning, an influential hypothesis has posited that tonic DA reflects average reward availability in a given context [34]. In other domains, however, tonic DA’s role has been best explained as reflecting precision within a Bayesian framework [52], which we discuss below. The view that DA modulates precision (operationalized as the signal-to-noise ratio) has empirical grounding in interval timing [53–57], motor control [58], vision [13, 14], and audition [59]. Interestingly, beyond *true* precision, DA has also been associated with *estimated* precision, or the agent’s estimate of its actual precision (or its confidence), independently of any changes in true precision [13, 60–62]. This is an important distinction, because a mismatch between true and estimated precision results in an underconfident or overconfident agent, which may affect behavior. In sum, the dual role of DA in true and estimated precision—and under what conditions this role holds—remains elusive. Once more, the duality of DA is not well understood.

Inspired by recent attempts to integrate the domains of reinforcement learning and interval timing [49–51], we will begin by introducing the concept of ‘rational inattention,’ borrowed from behavioral economics [63–65]. We will show that this framework unifies the two influential, yet seemingly distinct, algorithmic theories of tonic DA. We will then show that this framework predicts various empirical phenomena of reinforcement learning and interval timing, which cannot be explained by either theory alone.

## Results

### DA and rational inattention

To show how our account unites the two theories of tonic DA, let us begin by describing each one independently. First, under the average reward theory, tonic DA reports average reward availability in the current context. This theory has its roots in the observation that high tonic DA levels promote vigorous responses and high response rates in reinforcement learning tasks [34]. For a theoretical underpinning to this empirical phenomenon, Niv et al. [34] argued that animals in high-reward contexts should capitalize on the abundance of rewards with much of the same behaviors observed in hyperdopaminergic animals (high response vigor and high response rates). They thus proposed that tonic DA provides the link between average reward availability and the animal’s behavioral response, i.e., tonic DA levels report average reward. More concretely, in this view, DA can be thought of as reporting the opportunity cost of ‘sloth’, or the loss incurred by not capitalizing on the available rewards (in high-reward contexts, every passing moment not spent collecting reward is a wasted opportunity). Given that sloth is energetically inexpensive (and thus appealing), the average reward theory predicts that the animal will occupy this motivational state under low DA conditions (no use spending energy when reward potential is small), but will be increasingly incentivized to act quickly and vigorously as DA increases. The relationship of DA with responsivity and motivation has indeed been well documented [66–80].

Under the Bayesian theory, on the other hand, tonic DA signals the precision with which internal or external cues are stored and represented [52]. Thus under high DA, signaling high precision, the animal is more confident in its stored representations compared to the contextual information, and relies on them more heavily during decision making. This increased reliance can be probed by examining conditions under which the cue representations are put in conflict with other sources of information. Most commonly, this entails comparing the ‘bottom-up’ information (e.g., sensory cues and their stored representations) with ‘top-down’ information (e.g., prior beliefs about the cues, based on the context): Under high DA, the animal weights bottom-up information more heavily, whereas low DA promotes top-down information. In Bayesian terms, which we describe explicitly in the next section, DA increases the animal’s estimate of the likelihood precision relative to the prior precision. This theory has been used to explain behavioral aspects of DA-related pathologies such as autism [81–84], schizophrenia [85–87], and Parkinson’s disease [41].

In thinking about the Bayesian theory of DA, it is important to distinguish between the *estimated* precision (what the agent perceives its precision to be) and *true* precision (what its precision actually is). True precision increases through an increase in signal-to-noise ratios, which improves performance. On the other hand, an increase in estimated precision, without an equal increase in true precision (unwarranted confidence), can actually *impair* performance. Recent work has shown that, depending on the circumstance, DA can influence true precision, estimated precision, or both, such as in interval timing [41, 53–57], motor control [58], vision [13], and memory [88]. However, why DA would freely modulate estimated precision independently of true precision, in the first place, is not well understood. After all, under normal circumstances, precision miscalibration is maladaptive (see ‘Precision miscalibration’ section).

Each theory outlined above succeeds in explaining a subset of empirical findings, as we will show. But how can tonic DA reflect both average reward availability and precision? We unite these two theories of DA under the framework of rational inattention, inspired by ideas originally developed in economics [63–65]. Rational inattention posits that cognitive resources are costly, and therefore will only be spent when the agent is incentivized to do so. In particular, ‘attention’ to a stimulus is the cognitive process through which the agent reduces its uncertainty about the stimulus (see [89] for related definitions), and can be formalized in terms of precision: With increased precision, the agent will have greater certainty about its environment, which will increase its ability to accumulate rewards, but it will also incur a greater cognitive cost. Consider, then, a task in which an agent can improve its performance by attending to certain stimuli. As average reward availability increases, the agent will be increasingly incentivized to pay the cognitive cost necessary to increase precision. In this way, rational inattention provides the critical link between average reward and precision, and we hypothesize that this coupling is instantiated through DA: By reporting average reward, DA determines precision.

In the next sections, we will formalize this simple intuition and show how it allows us to expand the scope of experimental predictions made by each theory individually, while also conceptually connecting a number of seemingly distinct views of tonic DA.

### Bayesian inference

Before presenting the model, let us briefly discuss Bayesian inference, which will help us formalize the notion of precision (both true and estimated) and its effects on performance.

Suppose an animal is learning some parameter *μ*. For instance, *μ* may represent the expected reward obtained from some reward source, or the temporal interval between an action and an outcome. Because various stages of this learning process are characterized by noise [90], beginning with the nature of the parameter itself (stimuli are seldom deterministic) and leading up to the storage process (neurons are noisy), the learned information can be represented by a distribution over parameter values. This is referred to as a likelihood function (gray curves in Fig. 1; light and dark gray curves represent likelihood functions for a parameter with small and large magnitude, respectively). The spread of this distribution reflects the precision, or reliability, of the encoding process: High precision will lead to a tight distribution (Fig. 1A), whereas low precision will lead to a wide distribution (Fig. 1B). For simplicity, we take these distributions to be Gaussian throughout.

**Figure 1:**
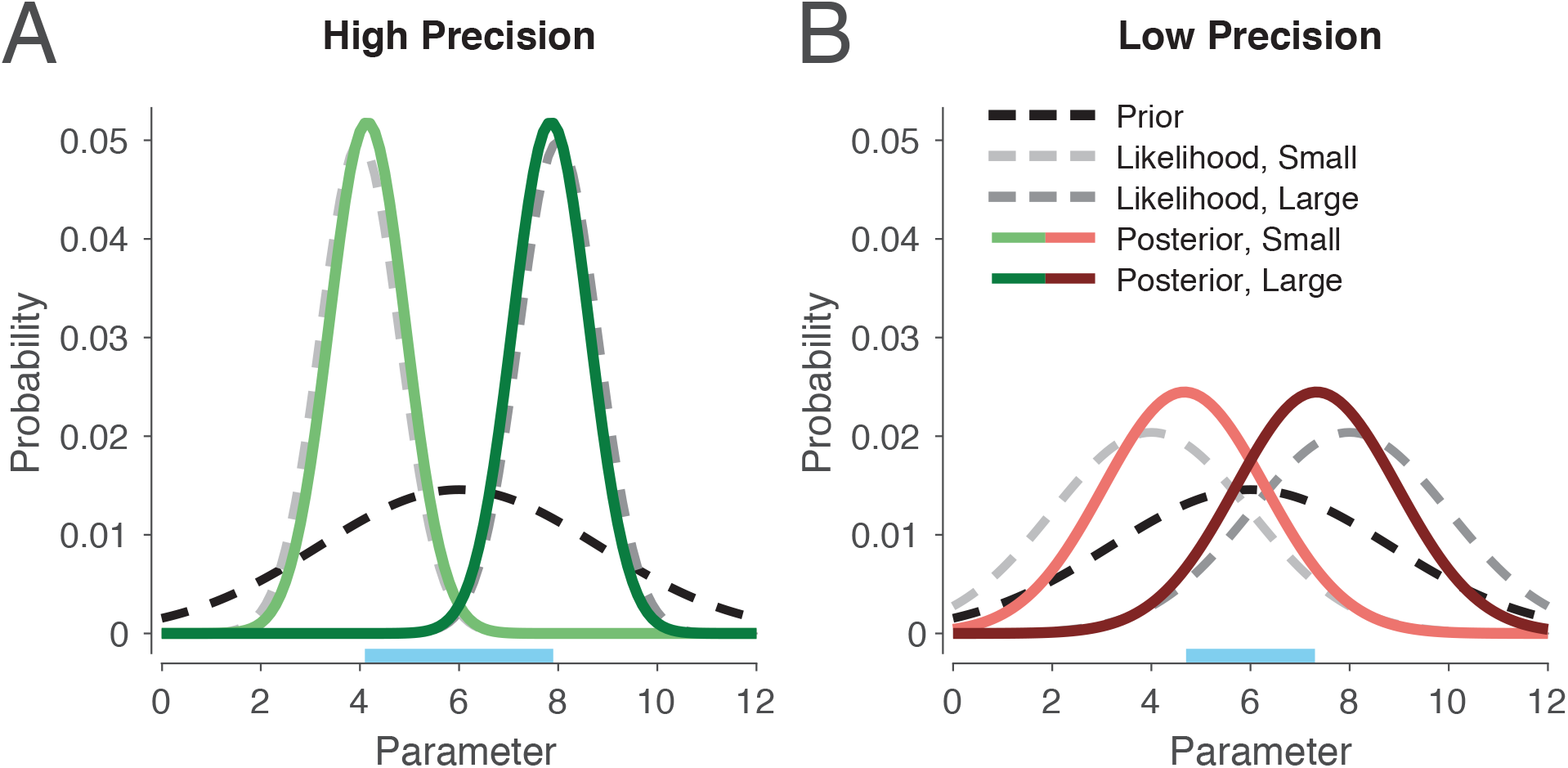
Illustration of Bayesian inference for a task with two parameters of different magnitudes, one small and one large, under either high or low precision. The posterior is the normalized product of the prior and likelihood. (A) When likelihood precision is high compared to prior precision, the posterior will remain close to the likelihood. (B) As the ratio of likelihood precision to prior precision decreases, the posterior migrates toward the prior. Note here that likelihood precision controls the distance between the posterior means (compare lengths of blue segments on the x-axis), a point we return to later.

During subsequent decision making, the animal must use this likelihood function to produce a learned estimate of *μ*, which we denote by 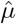. An optimal agent will use all the information available to it in order to produce this estimate, including its knowledge of the current context, or the ‘prior distribution’ (black curves in Fig. 1). For instance, if reward sources in this context tend to yield relatively larger rewards, or intervals tend to be longer, then it makes sense for the animal to produce an estimate that is slightly larger than the likelihood mean, especially when the precision of the likelihood is low (i.e., when it is not very reliable; compare black curves with light gray curves). Bayes’ rule formalizes this intuition [91–95] and states that an optimal agent will take the product of the likelihood and prior to compute the posterior distribution over parameter values (green and red curves in Fig. 1):

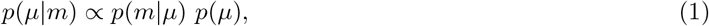

where *m* represents the stored (subjective) values, *p*(*m*|*μ*) is the likelihood, *p*(*μ*) is the prior, and *p*(*μ*|*m*) is the posterior. Under standard assumptions for Gaussian distributions [43, 95–97], the estimate 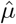 obtained by a Bayesian ideal observer will correspond to the posterior mean, which can be computed using the two quantities characterizing each distribution, their means and precisions:

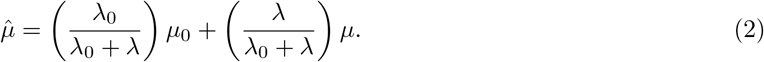

Here, *μ*_0_, *λ*_0_, *μ*, and *λ* represent the prior mean, prior precision, likelihood mean, and likelihood precision, respectively. In words, Eq. (2) states that the posterior mean 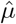 is a weighted average of the prior mean *μ*_0_ and the likelihood mean *μ*, and their respective precisions *λ*_0_ and *λ* constitute the weights after normalization. Hence, the tighter each distribution, the more it pulls the posterior mean in its direction (compare Fig. 1A and B). This is consistent with the intuition that one should rely more on what is more reliable. This type of optimal computation has been observed in many different domains [98].

In summary, when incoming information is noisy (low precision), an optimal agent will more strongly modulate its responses based on context. On the other hand, when the agent is very confident in incoming information (high precision), it will show little influence by context.

### Model description

Consider now a task where an animal must learn certain parameters *μ* in order to maximize its expected reward. For instance, in a temporal task, a rat may be trained to wait for a fixed period before pressing a lever in order to receive a drop of sucrose. Alternatively, in a reinforcement learning task, it may need to learn the magnitudes of rewards delivered from two different sources in order to choose the source with higher expected reward in the future. We can model these problems with the general encoding-decoding framework shown at the bottom of Fig. 2. Here, the animal transforms the objective stimulus *μ*—the duration of a timed interval or the magnitude of a reward—into a likelihood distribution *p*(*m*|*μ*) (encoding), which, during the performance stage, it then uses to produce its estimate 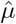 of the original stimulus via Bayes’ rule (decoding).

**Figure 2:**
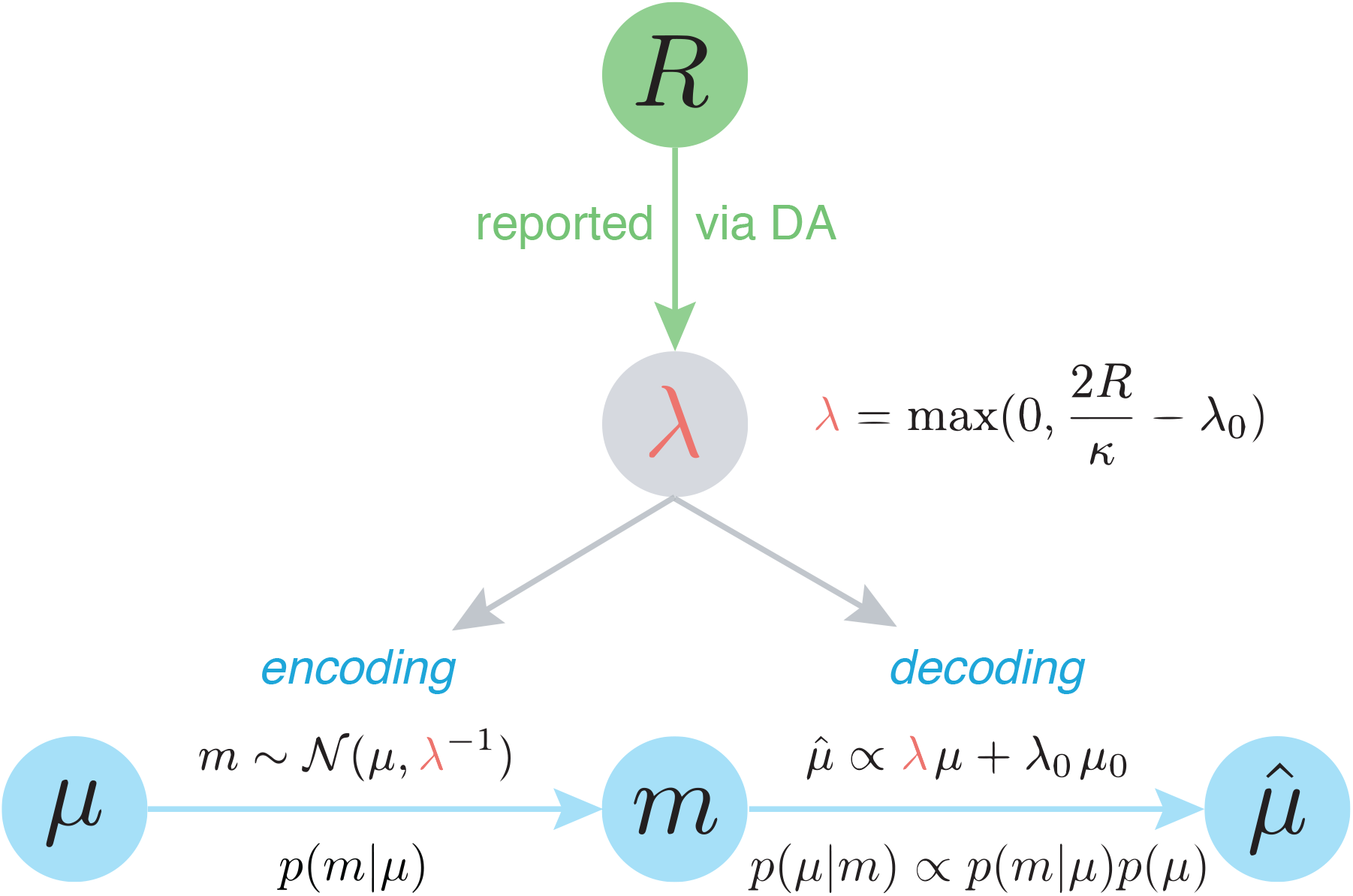
Rational inattention model of DA. Under rational inattention, average reward controls the likelihood precision through tonic DA. As derived in the next section and summarized in the top equation, increases in average reward increase likelihood precision, which in turn affects both encoding and decoding. Because average reward is a property of the context, DA can relay the likelihood precision (i.e., the precision of encoding) to the decoding stage, even when encoding and decoding are temporally decoupled. *R*: reward; *κ*: unit cost of attention (or information; see next section); *λ*: likelihood precision; *λ*_0_: prior precision; *μ*: likelihood mean (objective values); *m*: subjective (stored) values; *μ*_0_: prior mean; 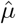: posterior mean (estimate of *μ*); *p*(*m*|*μ*): likelihood; *p*(*μ*): prior; *p*(*μ*|*m*): posterior.

This framework sheds light on the distinction between true and estimated precision. In the previous section, we implicitly assumed that the *encoding* precision *λ* is faithfully relayed to the *decoding* stage for Bayesian inference (i.e., we assumed the same *λ* for both stages). However, increasing *λ* selectively during decoding will increase the agent’s estimated, but not its true, precision. We refer to this as ‘precision miscalibration.’ Here, Bayesian inference will overestimate the precision with which encoding occurred, resulting in an excessive reliance on the likelihood compared to the prior (Fig. 1). Such miscalibration is maladaptive, and in a subsequent section we show that optimal performance occurs when the same precision used during encoding is also used during decoding.

Empirical evidence supporting the Bayesian theory suggests that DA influences both the true and estimated precision. But then, how is precision faithfully maintained across stages? After all, encoding and decoding can be temporally separated, and, given the natural fluctuations of tonic DA, what is the guarantee that the DA level would be the same at both stages?

We hypothesize that tonic DA reports the single quantity of average reward in a given context. According to rational inattention, this determines the precision *λ*. Imagine then that an animal experiences a high-reward context, learns its average reward, then leaves that context. If at a later date, the animal is returned to that same context, then its behavior there will depend on the average reward that it had previously learned, which is encoded in the DA signal. This means that the appropriate estimated precision can be retrieved from the DA signal when in that context. In this manner, by reporting average reward, DA during the encoding stage controls the true precision, and during the decoding stage, determines a faithful estimate of this true precision.

Of note, this soft ‘guarantee’ against precision miscalibration breaks down when DA levels are manipulated directly: For instance, while generally high levels of DA will increase both the agent’s true and its estimated precision, increasing DA selectively during decoding will increase the agent’s estimated, but not its true, precision. In the next sections, we will relate this miscalibration to experimental studies in which DA was acutely manipulated during the decoding stage. We will also show how this framework subsumes the view that DA implements ‘gain control’ on action values during action selection, without having to posit additional computational machinery.

In summary, by communicating the single quantity of context-specific average reward during both encoding and decoding, DA can achieve two seemingly distinct functions—controlling the precision of encoding incoming information as well as modulating the reliance on previously learned information when executing decisions.

### Mathematical description of rational inattention

Rational inattention views the agent as weighing the rewards gained from increased precision against the cost of this increase. With too much of an increase in precision, the cognitive cost may outweigh the additional rewards, whereas with too little of an increase in precision, the agent may forgo uncostly potential rewards. Let us begin by formalizing these two terms. This will then allow us to characterize the precision needed to maximize the agent’s utility (rewards minus cost).

In this section, we assume a perfectly calibrated agent, and hence make no distinction between estimated and true precision. The effects of precision miscalibration will be examined in the next section.

Consider an objective stimulus *μ*, such as a sensory cue or an environmental variable, that the agent must estimate. Let *x* denote the signal actually encoded by the agent, which is subject to noise that is reducible at a cost. Neurally, this may refer to noise at any level from signal transduction to storage, causing the stored neural representation *x* to be different from *μ*. We can model *x* as being drawn from a Gaussian distribution with mean *μ* and precision *λ* (the likelihood distribution), i.e., 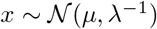, where the agent has control over *λ* (Fig. 2). The latent variable *μ* (the ‘source’) is drawn from some prior distribution representing the context, which we also take to be Gaussian for simplicity, with mean *μ*_0_ and precision *λ*_0_, i.e., 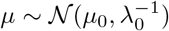. Then by Eq. (2), the posterior mean can be written as

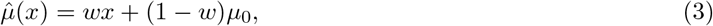

where

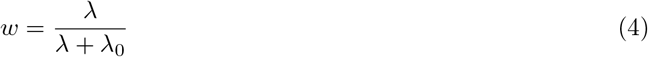

is the weight given to the likelihood mean, bounded by 0 and 1. We can now compute the average error in estimating the source. Intuitively, when *λ* is small, the estimate will migrate toward the prior mean, causing a large mismatch between the source and its estimate, whereas when *λ* is large, this migration—and hence the error—is small. For analytical convenience, we quantify this error using a quadratic error function:

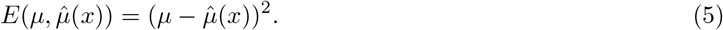

Using standard Gaussian identities, the ‘conditional’ error is given by

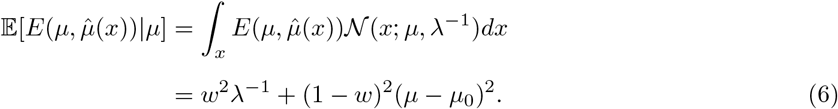

The conditional error can be understood as the agent’s expected response variance for a particular source. To compute the *overall* response variance (the ‘marginal’ error), we average across all sources:

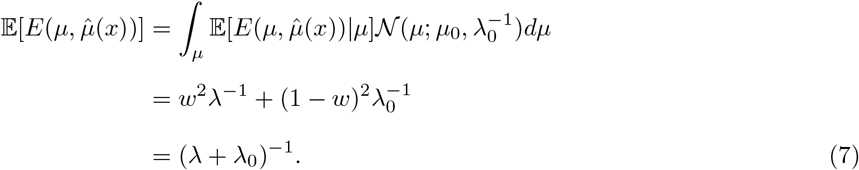

Thus an increase in the encoding precision *λ* decreases the marginal error, which in turn improves performance.

Next, let us formalize the cognitive cost of collecting information (reducing uncertainty) about the environment to improve performance. An analytically tractable choice of attentional cost function is the mutual information [99, 100], which measures the expected reduction in uncertainty due to the observation of the signal *x*:

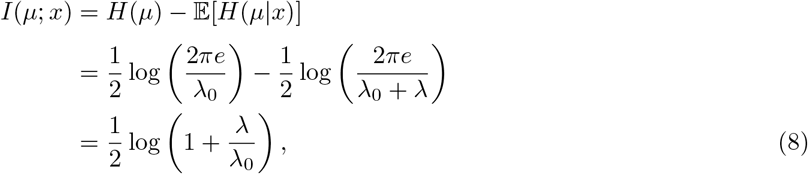

where *H*(*μ*) denotes the entropy of the probability distribution of *μ*. Intuitively, the uncertainty about *μ*, before observing the signal, can be quantified as the entropy of the prior distribution (high entropy, or uncertainty). On the other hand, after observing the signal, the uncertainty is represented by the entropy of the posterior distribution (lower entropy). The mutual information measures this reduction, which rational inattention assumes comes at a cost. The second equality follows from the entropy of Gaussian distributions with precision *λ*_0_ (the prior distribution; first term) and precision *λ*_0_ + *λ* (the posterior distribution; second term). Note that the posterior precision is simply the sum of the prior and likelihood precisions.

The choice of mutual information is appealing because it is tractable, displays the correct behaviors (increases in mutual information increase precision) [101], and is not restricted to a particular neural implementation. This final point is important given that the biological basis of cognitive effort remains unclear, despite attempts to link the two [102]. Indeed, while we assume the agent is primarily concerned with the cost of collecting and storing units of information, the rational inattention framework remains agnostic to the source of this cost at a biological level.

Assuming an increase in precision can improve performance, which for convenience we take to be linear with accumulated rewards, we can now combine the attentional cost function with the error function to define the attentional optimization problem:

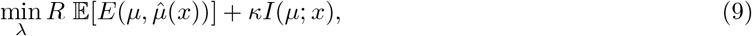

where *R* is the reward incentive, which we propose is reported by DA, and *κ* > 0 is the unit cost of information. As an incentive, *R* formally refers to the subjective, rather than the objective, value of the reward. This distinction will not be important for our model unless the two are specifically dissociated (e.g., through changes to satiety or baseline reward availability; see ‘DA and controllability’ section).

The agent seeks to minimize both the performance error and the (costly) reduction of uncertainty, which are weighted by the reward incentive and the unit cost of information, respectively. The idea here is that a unit increase in information decreases error (which leads to higher utility) but increases costs (which leads to lower utility). Thus if the agent pays *too* much attention to the task, the costs may outweigh the benefits, whereas if the agent pays *no* attention to the task, it may not reap any rewards. For our choice of cost and error functions, a middle ground exists that optimizes the agent’s utility (i.e., solves the optimization problem):

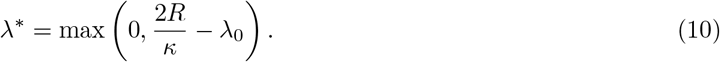

The rational inattention solution has three intuitive properties: First, attention to the signal increases with reward incentive (*R*). Second, attention to the signal decreases with the cost of information (*κ*). Third, attention to the signal decreases with prior precision (*λ*_0_). In other words, if the agent is more confident about the source before observing the signal, it will pay less attention to the signal. After all, there is no need to spend energy gathering information about a source when that source is already well known (Fig. S1).

We can also ask how the marginal error changes with respect to *κ* and *λ*_0_ by taking partial derivatives:

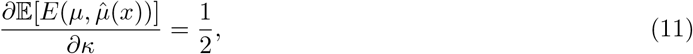

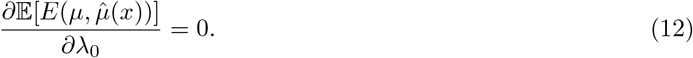

Thus, marginal error increases linearly with attentional cost, but does not vary with prior precision, since a unit increase in prior precision will be compensated for by a unit decrease in the optimal likelihood precision. Note also that the relationships are separable: The effect of attentional cost on marginal error does not depend on prior precision, and vice versa.

It is important to note that one can construct cost and error functions where such a middle ground is not attainable. For example, if the information cost and performance benefit take exactly equal but opposite shape, then the agent should always either increase its precision to infinity (if *R* > *κ*) or decrease it to zero (if *R < κ*). Our choice of functions, while principled, primarily serve to capture the intuition of the rational inattention framework.

Finally, we note that rate-distortion theory offers a normative interpretation of our optimization problem: If we accept that there is an upper bound on the bit rate of perception, then optimizing reward subject to this bit rate constraint will lead to Eq. (9) (the Lagrangian function), a standard result from rate-distortion theory [103].

### Precision miscalibration

Let us now analyze the effects of miscalibrated precision on the accuracy of a Bayesian agent. With *λ* and *λ′* denoting true and estimated precision, respectively, we set *c* = *λ′/λ*, which can be thought of as a miscalibration factor: When *c* < 1, precision is underestimated, and when *c* > 1, precision is overestimated. Here we examine the error incurred by miscalibration.

Following the derivations in the previous section, the signal weight can now be written as

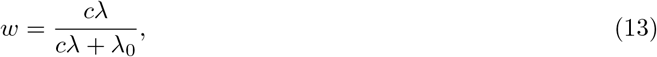

and the marginal error is

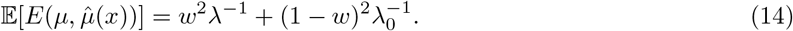

Taking the partial derivative of this expression with respect to *c* and setting it to 0, we find, consistent with intuition, that the error-minimizing miscalibration factor is *c* = 1. Thus, as an example, any experimental manipulation that increases estimated precision *λ′* without increasing true precision *λ* will produce a miscalibration factor greater than 1 and thereby incur an increase in error. Intuitively, a miscalibration factor of *c* > 1 means that both the signal *and* noise are amplified.

Note here that ‘miscalibration’ only refers to differences between true and estimated precision, and not to differences between the stimulus *μ* (or likelihood) and estimate 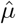 (or posterior). It is possible for *μ* and 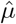 to be different without precision miscalibration, as shown in the previous section.

### Relationship with experimental data

#### DA and reinforcement learning

Let us now examine the predictions of our framework for the special case of reinforcement learning. Here, the task is to learn reward magnitudes and subsequently select the appropriate actions.

Our first prediction is that high DA during decoding will silence contextual influence, or, in Bayesian terms, decrease the influence of the prior. As illustrated in Fig. 1, an increase in likelihood precision, via high DA, amplifies the difference between the posterior means (compare blue horizontal segments on the x-axis). The parameter being reward magnitude here, this amplification makes the agent more likely to choose the action with the higher reward, i.e., to exploit. Similarly, low DA leads to low estimated precision and a significant Bayesian attraction toward the prior, which decreases the difference between the posterior means and promotes exploration (Fig. 3B). Indeed, previous work has suggested that DA controls the exploration-exploitation trade-off, whereby high DA encourages exploiting the option with the highest reward, and low DA encourages exploring other options [28–30] (but see [31] and Discussion). For instance, Cinotti et al. [30] trained rats on a non-stationary multi-armed bandit task with varying levels of DA blockade. The authors observed that the degree of win-shift behavior, representing the drive to explore rather than to exploit, increased with higher doses of the DA antagonist flupenthixol (Fig. 3A). Previous reinforcement learning theories of DA have explained this finding by suggesting that, during action selection, DA mediates gain control on action values [27, 104]. In the Methods, we show that DA’s decoding effect in our framework is equivalent to controlling the temperature parameter in the softmax function, a standard form of gain control [105, 106]. Thus, rational inattention endogenizes DA’s role in exploitation, without needing to posit a separate gain control mechanism for DA that only appears during performance.

**Figure 3:**
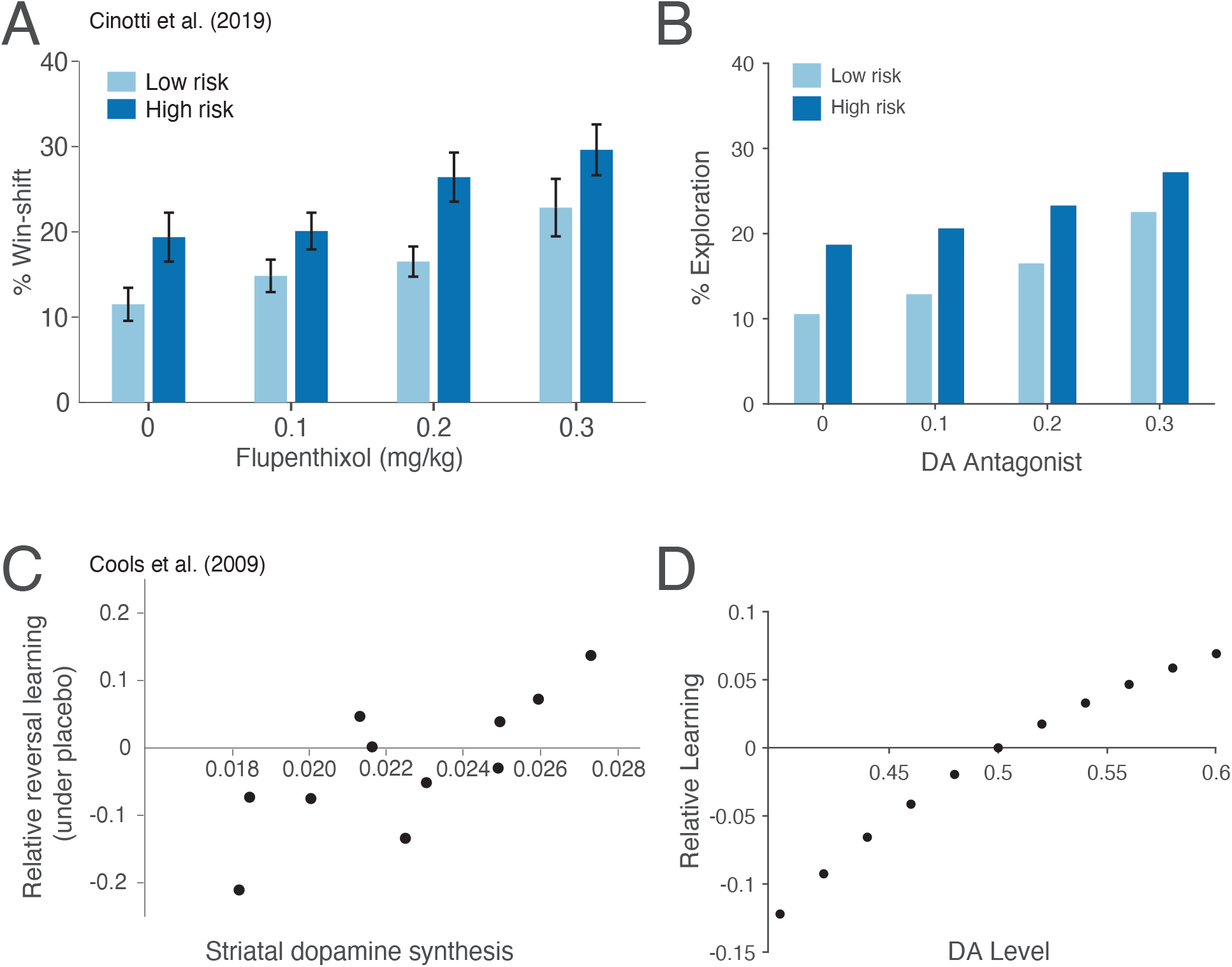
Rational inattention and reinforcement learning. (A) Using a non-stationary three-armed bandit task, Cinotti et al. [30] have shown that the DA antagonist flupenthixol promotes exploration (win-shift behavior). The task furthermore included two different risk levels: In low-risk conditions, one lever was rewarded with probability 7/8, and the other two with probability 1/16 each; in high-risk conditions, one lever was rewarded with probability 5/8, and the other two with probability 3/16 each. The effect of DA was evident for both conditions, and win-shift behavior was more pronounced in high-risk conditions across DA levels. Figure adapted from Cinotti et al. [30]. (B) Our model recapitulates these results: As DA decreases, likelihood precision decreases, which in turn reduces the difference in posterior means, and encourages exploration. The effect of risk on exploration follows from the reduced differences in likelihood means (and, by extension, posterior means) in high-risk compared to low-risk conditions. (C) Cools et al. [23] have shown that human subjects with high DA synthesis capacity learn better from unexpected rewards than from unexpected omissions of reward, whereas subjects with low DA synthesis capacity learn better from unexpected omissions than unexpected rewards. Relative accuracy is the accuracy following unexpected rewards minus the accuracy following unexpected punishments, taken to indicate the extent of learning from positive feedback compared to negative feedback. Figure adapted from Cools et al. [23]. (D) Our model recapitulates this result: High DA shifts the agent’s beliefs toward expecting positive feedback. Thus the speed of convergence for learning high rewards increases. Similarly, the speed of convergence for learning low rewards increases under low DA. The asymmetry in relative learning is due to an asymmetry in precision: Under high DA, likelihood precision is high, so the prior has a weaker effect than under low DA. For (B, D), see Methods for simulation details.

Our second prediction concerns the modulation of learning by DA. Learning can be thought of as iteratively weighing incoming information against a previously learned estimate to produce a new estimate. In so doing, the animal also learns the distribution of stimuli, which allows it to construct a prior for its context, as in Fig. 1. Under our framework, high DA signals high average reward in the context. Therefore, an agent under high DA should expect—and thus initialize its prior at—high rewards. This will result in faster learning of high rewards compared to low rewards, or equivalently, of positive feedback (rewards higher than expected) compared to negative feedback (rewards lower than expected). Similarly, under low DA, negative feedback will be learned better than positive feedback (Fig. 3D). Indeed, tonic DA levels have been shown to control the relative contribution of positive and negative feedback to learning [4, 5, 23, 107]. For instance, Cools et al. [23] repeatedly presented human subjects with a pair of stimuli (images), where one stimulus was associated with reward, and the other with punishment. On each trial, one of the stimuli was highlighted, and subjects had to predict whether that stimulus would lead to reward or punishment. Occasionally, unsignaled reversals of the stimulus-outcome contingencies would occur, so that the first stimulus to be highlighted after the reversal would result in an unexpected outcome. The same stimulus would then be highlighted on the subsequent trial. Accuracy on this second trial reflected the extent to which subjects learned from unexpected rewards (if the previously punished stimulus was highlighted) vs. unexpected punishments (if the previously rewarded stimulus was highlighted). The authors showed that subjects with higher DA synthesis capacity learned better from unexpected rewards, whereas those with lower DA synthesis capacity learned better from unexpected punishments (Fig. 3C). Interestingly, under rational inattention, learning better from positive or negative feedback is not a bias, but rather an optimal strategy.

It should be noted here that while striatal DA synthesis may in principle affect both phasic and tonic levels, the results of Cools et al. [23] cannot be explained as simply amplifying phasic DA, which putatively encodes reward prediction errors, without affecting tonic DA. For instance, recent work has shown that synthesis capacity and striatal prediction errors are in fact negatively correlated [108]. Furthermore, differential learning by positive vs. negative feedback has also been observed using pharmacological manipulations, which affect tonic DA directly [4].

#### DA and interval timing

We now examine our framework for the case of interval timing. We will focus on reproduction tasks, in which subjects must produce a previously learned interval under different manipulations, although our predictions will apply equally well to discrimination tasks, in which subjects respond differently to intervals of different lengths (e.g., responding ‘short’ or ‘long’ depending on whether a new interval is shorter or longer than a previously learned one). For each reproduction result below, we model its discrimination counterpart in Supporting Information.

Our first prediction is that while timing under high DA levels will be nearly veridical, timing under low DA levels will be susceptible to interfering temporal stimuli (strong migration toward the prior; Fig. 4B). Indeed, unmedicated Parkinson’s patients strongly display this effect, referred to as the central tendency. Here, reproducing durations of different lengths results in shorter intervals being overproduced and longer intervals being underproduced [39, 40], and veridical timing is rescued with DA repletion [39] (Fig. 4A; see also Fig. S3A,B). Shi et al. [41] have shown that these behaviors conform remarkably well to a Bayesian framework in which DA modulates the precision of the likelihood. Rational inattention takes this one step further: Because DA reflects average reward, our framework also predicts the central tendency under low average reward conditions or satiated states (in which rewards are devalued). This is consistent with empirical studies that manipulated average reward and satiety through prefeeding [109–111], although motivation is a confound in these experiments (Fig. S4A,B).

**Figure 4:**
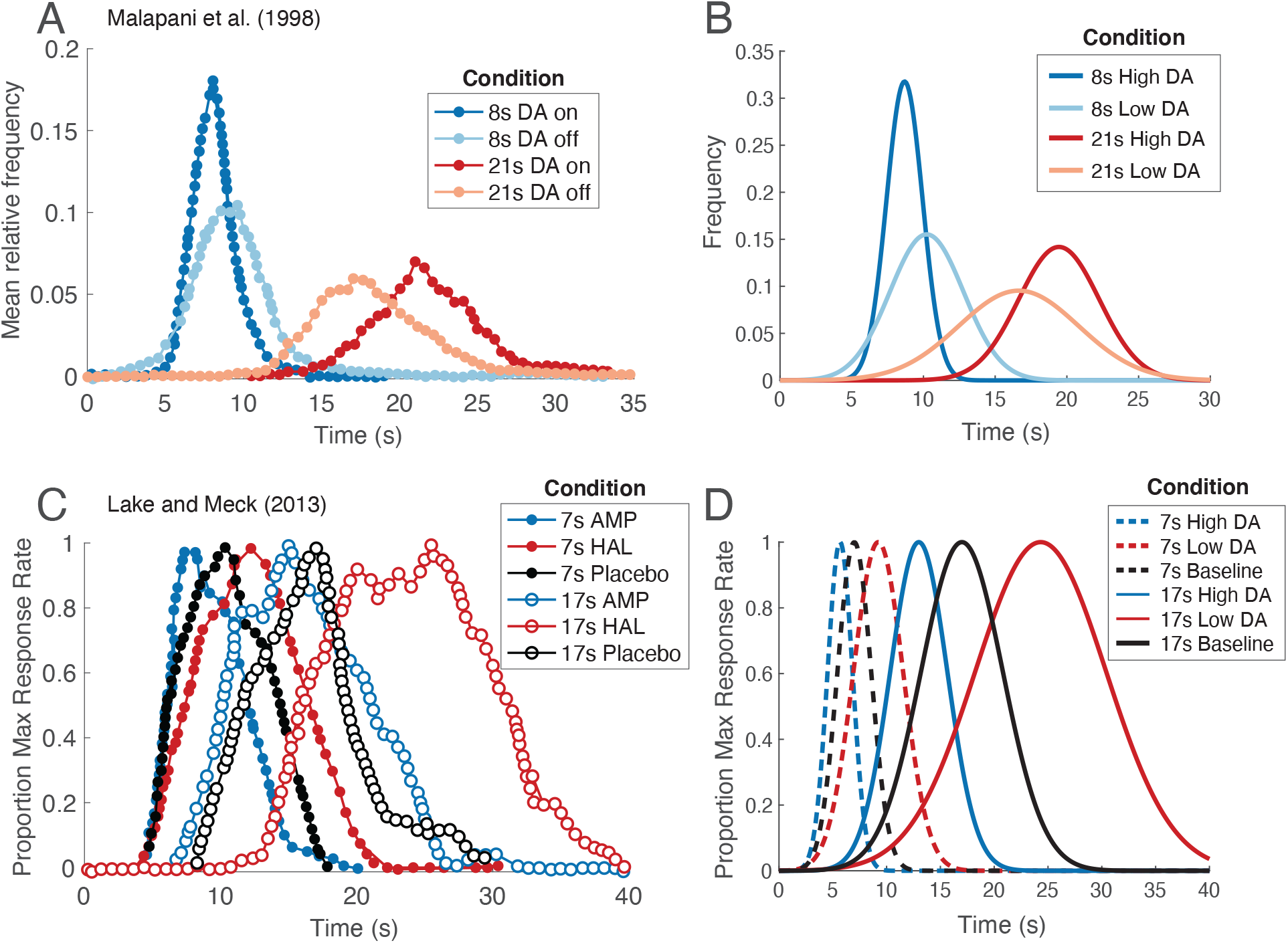
Rational inattention and interval timing. (A) DA masks the central tendency effect. Malapani et al. [39] have shown that when unmedicated Parkinson’s patients learn intervals of different durations in an interleaved manner, they overproduce the shorter intervals and underproduce the longer ones (light blue and orange curves). Medication rescues veridical timing (dark blue and red curves). Figure adapted from Malapani et al. [39]. (B) Our model recapitulates this effect: When DA is high, likelihood precision is high, and the posterior closely resembles the likelihood. When DA is low, likelihood precision is low, and the posterior migrates toward the prior. (C) DA increases the speed of the internal clock. Lake and Meck [17] trained healthy human subjects on reproducing a 7- or 17-second interval. They then acutely administered either amphetamine (DA agonist) or haloperidol (DA antagonist), and observed temporal reproduction that was consistent with either a faster or a slower clock, respectively. Note that the central tendency is not captured in this experiment because each of the 7-second and 17-second intervals was presented separately in a blocked manner, but plotted in the same figure for convenience. Figure adapted from Lake and Meck [17]. (D) Our model recapitulates this effect: When DA is high, temporal receptive fields must compress against objective time to increase likelihood precision. This results in a faster internal clock. When DA is low, temporal receptive fields expand to decrease likelihood precision. This results in a slower internal clock. For (B, D), see Methods for simulation details.

Our second prediction concerns acute manipulations of DA during decoding. A large body of work has shown that tonic DA affects interval timing by modulating the speed of the internal clock (or the ‘subjective time’), whereby higher DA levels lead to a faster clock [15–17, 37, 38, 112–116] (Fig. 4C). This finding has been replicated under different experimental paradigms to control for potential confounds (e.g., motivation; Fig. S3C,D). Effects on clock speed are also well documented in the behavioral literature: Here, the speed of the clock increases when animals are placed in high reward-rate contexts and decreases in low reward-rate contexts [117–123] (Fig. S4C). Clock speed similarly decreases in satiated states (i.e., in reward devaluation) [124, 125].

Our framework predicts these findings under the assumption that temporal precision is controlled by the internal clock speed. Recent empirical findings provide some support for this assumption. Indeed, a number of studies have identified ‘time cells’ (e.g., in striatum and medial frontal cortex [126, 127]), which seem to function as an internal timing mechanism: Time cells fire sequentially over the course of a timed interval, they tile the interval, and their activations correlate with timing behavior. In other words, their activations seem to reflect ‘temporal receptive fields.’ By definition, temporal precision is inversely related to the temporal receptive field width. Thus, any rescaling of the time cells will modify the precision but also change the speed of the internal clock (or change the mapping between objective and subjective time; Fig. S2). Rescaling of time cell activations has indeed been well documented [126, 127].

Given that the clock speed can change, how is it then that timing can ever be reliable? Under rational inattention, the answer is straightforward: Reporting context-specific average reward, DA maintains the same precision across both encoding and decoding. If precision is implemented through changes in the speed of the internal clock, it follows that the clock speed will also be the same during encoding and decoding, and, in general, timing will be reliable.

Let us now turn to acute manipulations of DA. Under rational inattention, an acute increase in DA levels at decoding increases estimated precision, implemented here by compressing the time cell activations against objective time (Fig. S2). This increases the speed of the internal clock, which results in underproduction of previously learned intervals. Similarly, acutely decreasing DA levels at decoding will slow down the internal clock, resulting in overproduction of stored intervals, consistent with the empirical findings (Fig. 4D). This framework also predicts the influence of average reward on clock speed (Fig. S4D) as well as that of different motivational states—Under rational inattention, reward and DA manipulations are equivalent.

Finally, note that the ability to keep track of time worsens as the interval duration increases. This worsening is known as Weber’s law, which asserts that the inverse square root of precision—or the standard deviation— increases linearly with time [128–130]. The predictions specific to our model do not depend on Weber’s law, whose underlying cause is still a subject of active debate [e.g., 129, 131, 132] (only the pattern of wider probability distributions for larger intervals depends on Weber’s law). However, for a completely determined mathematical model, we propose a rational-inattention-based derivation of this phenomenon in Supporting Information. This will infuse our model with quantitatively precise predictions without affecting our qualitative results.

#### DA and controllability

Finally, rational inattention makes the counterintuitive prediction that the effects of average reward in increasing the speed of the clock will be conditional on controllability. In other words, attentional cost is only paid if outcomes can be improved. Otherwise, there is no use in spending cognitive resources on a task beyond the animal’s control.

Bizo and White [133, 134] sought to directly examine the effect of freely delivered reinforcers on interval timing. To do so, they trained pigeons on a free-operant task that allowed for left-key responding and right-key responding. In each 50-second trial, only the left-key response was rewarded during the first 25 seconds, and only the right-key response was rewarded during the last 25 seconds. As expected, trained pigeons were more likely to select the left-key response early on, and the right-key later on. This response was quantified with a psychometric function, in which the probability of making a right-key response was plotted against time, and which took a sigmoidal shape. To examine the effect of free rewards, the authors additionally included a center key that would freely deliver rewards independent of the timing task. From the psychometric function, the authors fit a computational model of timekeeping to deduce a ‘pacemaker period’ (inverse of clock speed). Intuitively, a shift in the psychometric function to the left (being more likely to select the right-key response early) was taken to indicate that the clock speed increased (smaller pacemaker period), whereas a shift to the right was taken to indicate that the clock speed decreased (larger pacemaker period). The authors found that the speed of the internal clock varied *inversely* with freely delivered rewards (Fig. 5A), consistent with the principle of rational inattention: As freely delivered rewards became more available, the incentive to pay attention to the timing task decreased, causing precision, and thus clock speed, to decrease (Fig. 5B; Methods). On the other hand, when performance-contingent rewards became more available compared to freely delivered rewards, the clock speed increased (Fig. 5A,C). Experiments in mice suggest that DA levels are higher in tasks with controllable outcomes and lower under learned helplessness (lack of control) [135, 136], as predicted by our framework.

**Figure 5:**
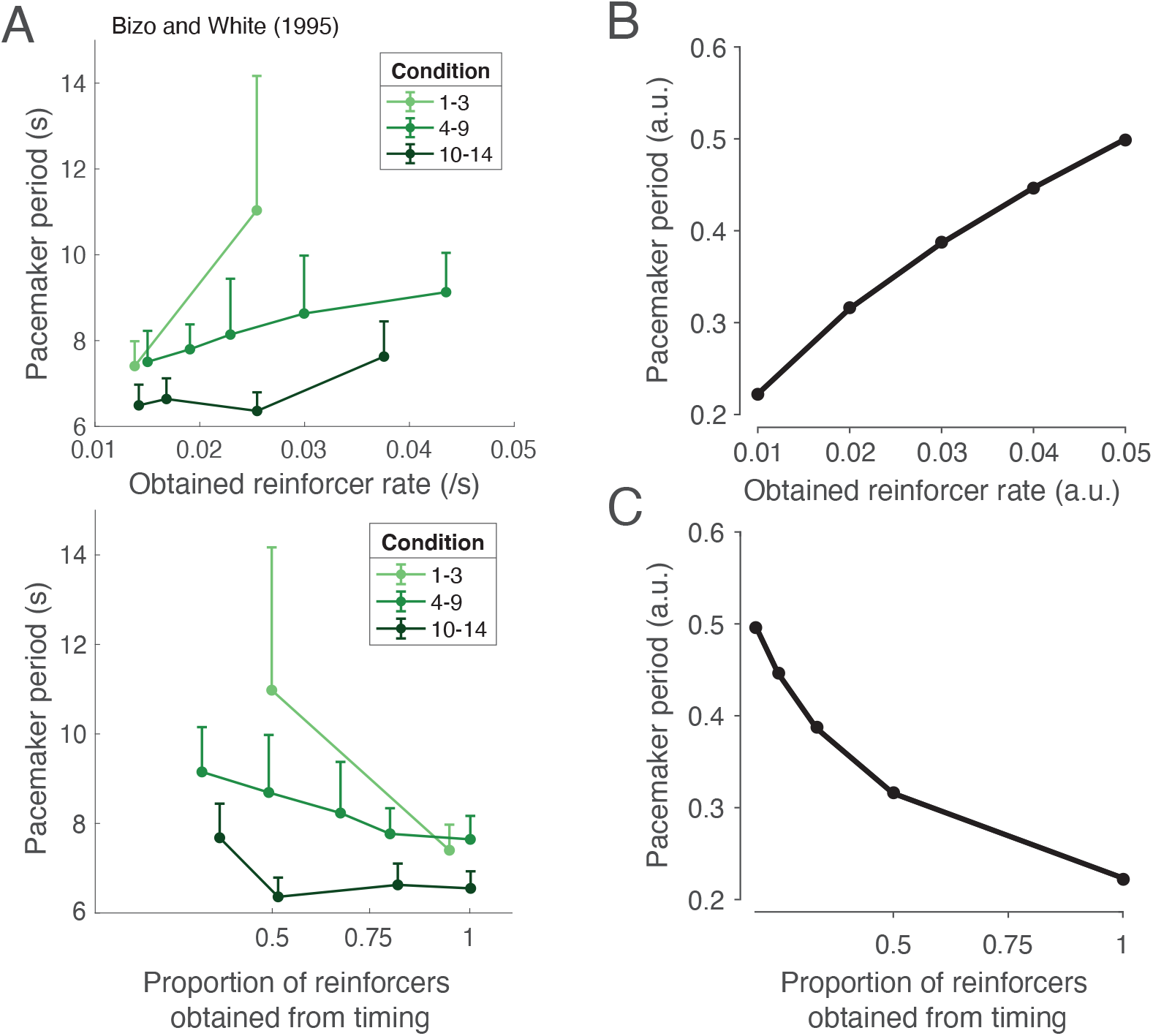
Rational inattention and controllability. (A) Controllability increases clock speed. Top panel: Bizo and White [134] have shown that when reinforcers are freely delivered, clock speed decreases (plotted here is the pacemaker period, or the inverse of clock speed, a parameter in their computational model which was fit to the empirical data). Bottom panel: On the other hand, when obtaining reinforcers is contingent on adequate timekeeping, clock speed increases. Light green, green, and dark green denote conditions in which free rewards were delivered during the intertrial interval, during both the intertrial interval and the trial, and during the trial only. We do not make a distinction among these conditions in our model. Figure adapted from Bizo and White [134]. (B, C) Our model recapitulates this effect: Under rational inattention, high average reward should only increase precision when this increase improves the ability to obtain rewards. (B) As free rewards increase and the added incentive of timing-contingent rewards decreases, clock speed will decrease. (C) On the other hand, as timing-contingent rewards increase, clock speed will increase. See Methods for simulation details.

Should the clock speed decrease to zero in the complete absence of controllability? It is reasonable to assume here that, even in this case, the animal should still pay *some* attention to the task, given the potential usefulness of observational learning for future performance. Additionally, in the real world, tasks overlap (a predator preparing to pounce uses the same visual information to identify its prey, assess its fitness, localize it well, and predict its next move), so reducing attention in a single subtask to zero without affecting the others is often not feasible.

Rational inattention also reconciles a longstanding question on temporal discounting and post-reward delays in animals: A large body of work has used intertemporal choice tasks to study the impact of delays on reward preferences in animals. In these studies, animals are given the choice between a small reward delivered soon, or a large reward delivered later. Choosing the smaller (sooner) reward has been taken to indicate a discounting of future rewards, with the extent of discounting often taken to reflect qualities like impulsivity and self-control. A closer look, however, has suggested that animals’ behaviors are consistent with an underestimation of time during the post-reward delay period, which is typically included after choosing the small reward in order to control for total trial durations and average reward rates [137–139]. Thus the apparent temporal discounting may simply arise from this underestimation of time during the post-reward delay, independent of reward value. For instance, Blanchard et al. [139] tested monkeys on an intertemporal choice task in which the post-reward delay was varied (Fig. 6A). By motivating and fitting a model in which the animals maximized long-term reward rate, they showed that the animals systematically underestimated the post-reward delays (Fig. 6B). The cause of this underestimation is still an open question [139]. However, rational inattention predicts this effect, as animals have more control over the outcome of the task before reward presentation than after it. Thus our framework predicts a slower clock during the post-reward delay, and behaviors consistent with an underestimation of time (Fig. 6C).

**Figure 6:**
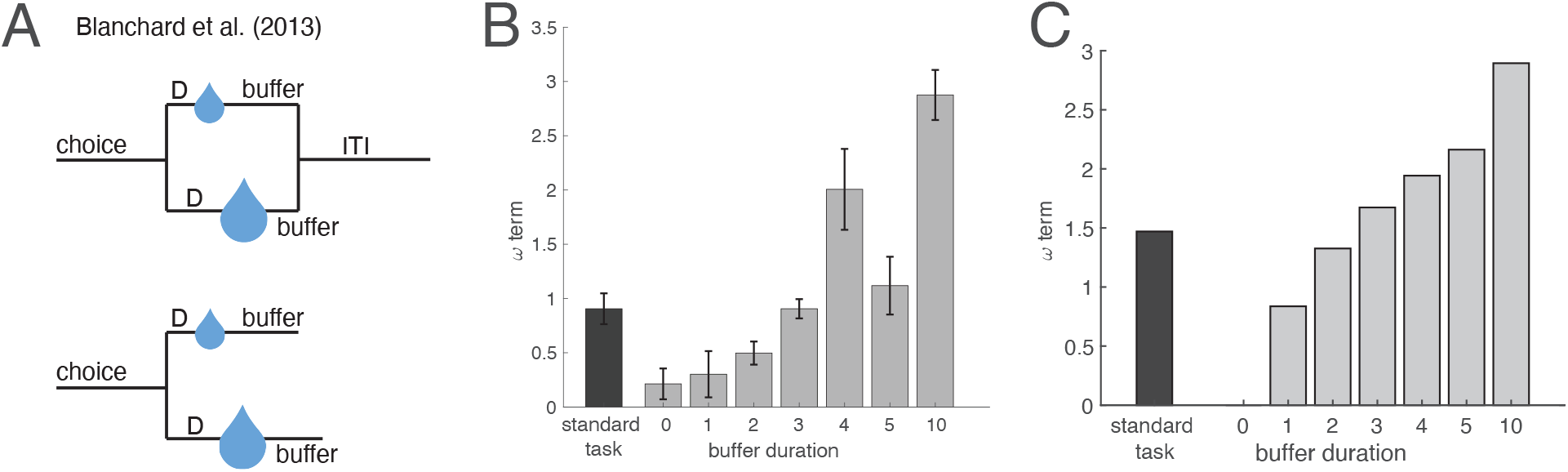
Rational inattention and post-reward delays. (A) Blanchard et al. [139] trained monkeys on an intertemporal choice task involving a small reward delivered soon or a large reward delivered later. Top: In the standard task, the total trial duration, or the sum of pre-reward delay (‘D’) and post-reward delay (‘buffer’), was fixed to 6 seconds. The average buffer duration was 3 seconds. Bottom: In the constant buffer task, the post-reward delay was fixed regardless of choice, and was either 0, 1, 2, 3, 4, 5, or 10 seconds. iti: intertrial interval. (B) Monkeys underestimate post-reward delays. By fitting monkey behavior to a model in which animals maximize long-term reward rate, the authors showed that the model fit value for the subjective estimate of post-reward delay (the ‘w term,’ described in the Methods) is smaller than its true value (buffer duration). This relationship held for the standard task and across all buffer durations in the constant buffer task. For (A, B), figures adapted from Blanchard et al. [139]. (C) Our model recapitulates this effect: Under rational inattention, precision increases during the period when outcomes can be more strongly controlled (i.e., the pre-reward delay), and decreases otherwise (i.e., the post-reward delay, before the subsequent trial begins). This creates the apparent effect of an underestimation of post-reward delays. See Methods for computational model and simulation details.

This result makes the untested prediction that optogenetically stimulating DA neurons during the post-reward delay should make animals less impulsive. Note here that the animal’s behavior will appear to be impulsive with a *faster* clock in the *pre-*reward delay, or a *slower* clock in the *post-*reward delay. This is because the larger/later option has a longer pre-reward delay, but the smaller/sooner option has a longer post-reward delay. Thus a faster clock during the pre-reward delay will disproportionately increase the perceived total trial length of the larger/later option (more impulsivity), whereas a faster clock during the post-reward delay will disproportionately increase the perceived total trial length of the smaller/sooner option (less impulsivity).

It is important to note that the discounting-free model does not invalidate reward discounting in general. While the intertemporal choice task in animals involves training over many trials, discounting in humans typically involves mental simulations (“Do you prefer $1 today or $10 in one month?”) that may involve neither an experienced pre-reward delay nor a post-reward delay. Nonetheless, humans systematically discount future rewards in these tasks [140].

#### Experimental predictions

The rational inattention framework makes a number of novel experimental predictions. Broadly, these can be classified based on any of three experimental manipulations (controllability, average reward, and DA level), two time courses (chronic and acute), two domains (reinforcement learning and interval timing), and two readouts (DA and behavior). Let us illustrate these with the testable example of controllability in reinforcement learning.

Consider a reinforcement learning task in which animals or humans are trained on a two-armed bandit, where arm *A* yields a small reward and arm *B* yields a large reward. In Experiment 1, subjects can sample the arms freely (high controllability), and the arms are denoted by *A*_1_ and *B*_1_. In Experiment 2, subjects are merely observers, so the arms are sampled for them (no controllability), and the arms are denoted by *A*_2_ and *B*_2_. Arms *A*_1_ and *A*_2_ yield the same reward, but each is accompanied by a distinct stimulus (e.g., sound); similarly for arms *B*_1_ and *B*_2_. After training on Experiments 1 and 2 separately, subjects are tested on a two-armed bandit consisting of arms *A*_1_ and *A*_2_ or arms *B*_1_ and *B*_2_ (each with its accompanying stimulus, so that the arms are distinguishable). The rational inattention framework predicts that, because of the central tendency, *A*_2_ will be more likely to be selected than *A*_1_, whereas *B*_1_ will be more likely to be selected than *B*_2_ (Fig. 7A). This effect cannot be explained by a preference for one type of controllability over the other: For B, the option trained under a *more* controllable context is preferred, whereas for A, the option trained under a *less* controllable context is preferred. Similarly, the effect cannot be explained by assuming better learning in one experiment over the other. Finally, this setup can control for any differences in sampling between the two experiments, as the choices in Experiment 2, which are made by a computer, can exactly match those made by the subject in Experiment 1.

**Figure 7:**
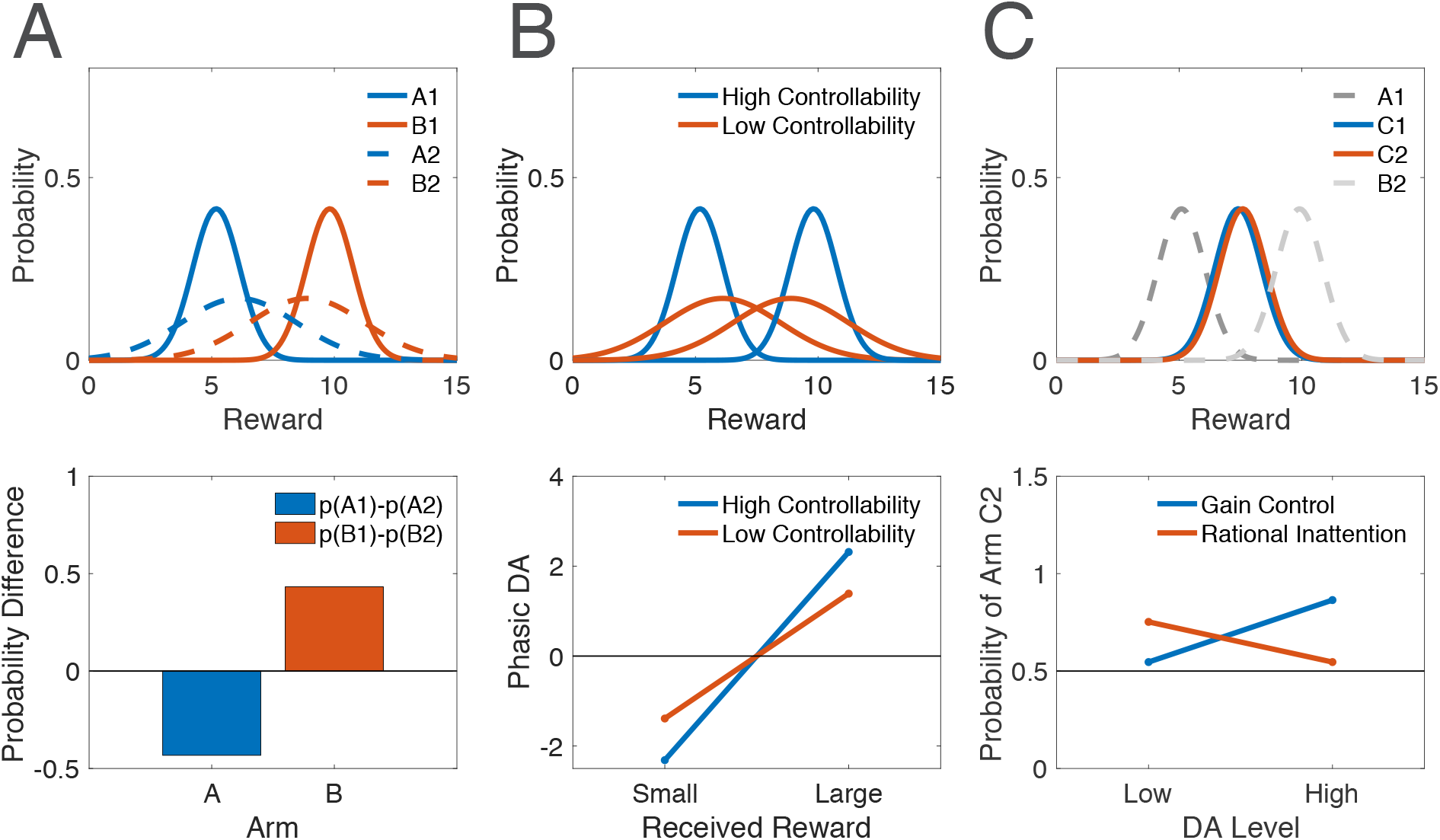
Experimental predictions of rational inattention. (A) Top panel: The difference in value estimates for two arms is higher under high controllability (solid curves) than low controllability (dashed curves). Bottom panel: After learning under each condition, the two arms yielding small rewards are compared. Rational inattention predicts that the arm trained under low controllability will be selected more often. On the other hand, when the two arms yielding large rewards are compared, that trained under high controllability will be selected more often. p(A1): probability of selecting arm A1; similarly for B1, A2, and B2. (B) Top panel: When a single arm yields a small or large reward with equal probability, the estimated deviation of actual outcomes from the mean reward will have larger magnitude under high-controllability learning than under low-controllability learning. Bottom panel: Thus the small reward will elicit a more negative phasic DA response, and the large reward will elicit a more positive phasic DA response, under high controllability than under low controllability. (C) Top panel: Arms C1 and C2 are identical, but C1 is trained with an arm yielding a smaller reward (A1), and C2 is trained with an arm yielding a larger reward (B2). The estimates for the two identical arms will be on opposite sides of their true value due to the central tendency. Bottom panel: After training, arms C1 and C2 are combined into a new two-armed bandit task, occurring under either high or low DA. The gain control hypothesis of DA predicts that the difference in their estimates will be amplified under high DA, thus making selection of C2 more likely than under low DA. On the other hand, rational inattention predicts that the central tendency will be reduced under high DA, which, in this task, will cause the two estimates to migrate closer to their true reward value (and therefore, to each other), in turn making selection of C2 less likely than under low DA. See Methods for simulation details.

Analogs of this experiment can be performed while varying average reward or DA levels instead of controllability. In these cases, the probability of selecting each action during training can further be compared across experiments: The probability difference in choosing each arm will be greater in Experiment 1 (high average reward or high DA) than in Experiment 2 (low average reward or low DA) due to the central tendency. This is similar to the experiment by Cinotti et al. [30] discussed previously, and can be interpreted as the animal exploiting under high average reward and high DA, and exploring under low average reward and low DA. (Note here that differences in sampling frequency for each arm are not controlled for.)

A second experiment can test value estimates by directly measuring *phasic* DA responses, which putatively report ‘reward prediction errors’ (RPEs), or the difference between received and expected rewards [1–3, 21, 22]. Consider then a task in which animals or humans are again trained on a two-armed bandit, either with (Experiment 1) or without (Experiment 2) controllability, but where one arm is stochastic. In particular, let arm *A* yield a reward that is either small or large, with equal probability. After some learning, when arm *A* is chosen, the large reward will elicit a phasic DA burst (received reward is larger than expected), whereas the small reward will elicit a dip (received reward is smaller than expected). This will occur in both Experiments 1 and 2. However, the phasic burst following *A*_1_ when a large reward is received will be greater than that following *A*_2_ (we use the same subscript notation as above). On the other hand, the dip following *A*_1_ when a small reward is received will be deeper than that following *A*_2_, again due to the central tendency (Fig. 7B). As above, this effect cannot be explained by a preference for one type of controllability or preferential learning in one experiment over the other. Also as above, analogs of this experiment can be performed by replacing controllability with average reward or tonic DA levels.

Third, using the same principles, we can design an experiment to distinguish between the gain control hypothesis of tonic DA and the rational inattention framework. Recall that, to explain tonic DA’s role in performance (as opposed to its role in learning), a number of authors have posited that tonic DA implements some form of ‘gain control’ on action values, whereby differences in action values between different options are amplified by high tonic DA levels, which promotes exploitation of the most rewarding option. The rational inattention framework subsumed this idea by showing that controlling precision mimics gain control, at least in simple cases (Methods). Let us then design an experiment to distinguish between these alternatives: Consider a two-armed bandit task of small and medium reward (Experiment 1) and medium and large reward (Experiment 2). After training on these pairs independently, the two identical (or nearly identical) medium rewards—one from each experiment—are combined into a new pair. The subject must make selections in this new two-armed bandit task, under either high or low DA. Rational inattention predicts that, because of the central tendency, the medium reward from Experiment 2 will be selected more often under low DA, with a smaller (or absent) effect under high DA. But this is the exact opposite prediction of the gain control hypothesis, whereby any differences in selection should be amplified under high DA (Fig. 7C). As a proof of concept, this result may begin to explain why some studies have found exploration to be enhanced under high DA [31], which thus far has been viewed as incompatible with the seemingly competing literature discussed above.

## Discussion

Questions on the roles of tonic DA abound. While two theories have put forth compelling arguments attributing tonic DA to either precision or average reward, it has remained unclear conceptually whether and how these two quantities are related to each other. Furthermore, within each domain, questions arise. Under the precision viewpoint, why would fluctuations in tonic DA separately influence both true and estimated precision, a seemingly suboptimal strategy when encoding and decoding are temporally decoupled? In reinforcement learning models, how and why does tonic DA implement gain control on action values during performance, and how does this relate to its role in learning? Rational inattention resolves these questions: By reporting the single quantity of context-specific average reward, DA first determines the precision with which encoding occurs, and second, faithfully relays the precision used during encoding to the decoding stage. This view unifies the two theories, while simultaneously endogenizing the ‘gain control’ function of DA and safeguarding against the suboptimality of precision miscalibration. Beyond DA, this framework takes an additional step toward integrating theories of reinforcement learning and interval timing.

In reinforcement learning, the rational inattention framework predicts that learning from positive and negative feedback is enhanced under high and low DA, respectively: Because DA signals average reward, an animal under high DA expects high rewards in these tasks. Thus under high DA, positive feedback is expected, and therefore more readily learned, whereas under low DA, negative feedback is more readily learned. Second, under rational inattention, high DA suppresses the contribution of context to the final estimate. Thus when two reward magnitudes are learned and subsequently compared, this suppression of interfering signals increases the difference between the estimated magnitudes. This in turn tips the exploration-exploitation balance toward exploitation of the higher reward.

In interval timing, rational inattention predicts that both high DA levels and high average reward result in a faster internal clock: Constrained by time cell rescaling, increases in precision occur through a compression of subjective time against objective time, and thus lead to a faster internal clock. We take these scalable time cells to constitute the internal timing mechanism [126, 127], which may be well-adapted to rapid learning of the temporal properties of an environment [141]. Similarly, low DA levels and low average reward lead to a slower clock. Low DA simultaneously increases the relative contribution of context (other signals) to the temporal estimate, which exaggerates the central tendency effect. Finally, rational inattention also predicts that the modulation of precision by average reward should only apply when obtaining the rewards is contingent on the agent’s performance. When rewards are freely delivered, precision should not increase, as it would be wasteful to spend cognitive resources on the present task. Faced with enough rewards that are not dependent on performance, the clock should instead slow down. The effect of controllability also implies that animals will underestimate the duration in a task that follows reward delivery but precedes the end of each trial, a long-known but poorly understood phenomenon of post-reward delays.

Rational inattention captures empirical phenomena that cannot be captured by either the average reward theory or the Bayesian theory alone (Table 1). While the average reward theory succeeds in explaining the effect of learning from positive vs. negative feedback, it fails in capturing all other presented phenomena. The Bayesian model is more successful, and can be extended to capture the exploration-exploitation findings in reinforcement learning as well as the DA manipulations in interval timing (central tendency and clock speed), but fails in capturing the effect of learning. Both theories fail in predicting the effect of average reward on the speed of the internal clock, the effect of controllability, and post-reward delays. In other words, the individual theories fail when the experimental manipulations and measured outcomes correspond to variables upstream and downstream of DA, respectively (i.e., nodes above and below the green arrow in Fig. 2, inclusive). When both are upstream of DA (thus involving controllability, average reward, and DA), the average reward theory is successful; when both are downstream of DA (thus involving DA and precision), the Bayesian theory is successful. As an example, consider the effect of controllability on the speed of the internal clock. The average reward theory predicts that animals should act more vigorously and with higher response rates when in high average-reward environments, which applies to timing-contingent and timing-noncontingent rewards alike. However, without some additional assumption, the theory does not predict that clock speed should increase in the first case and decrease in the second. Similarly, while the Bayesian theory can accommodate changes in clock speed, it lacks a theoretical foundation for why controllability should influence the clock (or even, as an intermediate step, the DA level).

**Table 1:** Summary of predicted phenomena by theory. The average reward theory is successful when controllability, DA, or average reward is manipulated, and DA or the reward estimate is measured. The Bayesian theory is successful when DA is manipulated, and behavior is measured. The rational inattention framework is successful with any combination of these variables.

|  |  | Average reward theory | Bayesian theory | Rational inattention |
| --- | --- | --- | --- | --- |
| **Reinforcement learning** | DA and learning from positive vs negative feedback | ✓ | ✗ | ✓ |
|  | DA and exploitation | ✗ | ✓ | ✓ |
| **Interval timing** | DA and the central tendency | ✗ | ✓ | ✓ |
|  | DA and clock speed | ✗ | ✓ | ✓ |
|  | Average reward and clock speed | ✗ | ✗ | ✓ |
| **Controllability** | Clock speed | ✗ | ✗ | ✓ |
|  | Post-reward delays | ✗ | ✗ | ✓ |

The framework we have presented is related to the hypothesis that DA in some tasks mediates the ‘cost of control.’ Notably, recent work by Manohar et al. [142, 143] has shown that subjects break the speed-accuracy trade-off in a saccadic selection task when motivated by reward, simultaneously increasing their speed and improving their accuracy. The authors propose a cost of control, which can be overcome with rewards, an effect they hypothesize is mediated by DA. While our work generalizes this result at the normative and algorithmic levels, the question of how the precision-cost trade-off is implemented neurobiologically remains to be determined in future work. It should be emphasized here that our framework removes the confounding speed-accuracy trade-off element, because in the tasks we model, agents cannot earn more reward by responding more quickly. For instance, Otto and Daw [144] have found that, when subjects are given a set amount of time to sequentially collect as many rewards as they want, they show higher error rates. This makes sense from a normative perspective: Spending too much time on one trial forces subjects to forgo future rewards. So under high reward rates, subjects should act faster, even if it means a higher error rate, because the total number of accumulated rewards will go up.

It is important to note that the story of DA and performance is not as linear as we have assumed so far. Rather, a significant body of work has shown that when very high DA levels are reached, an inverse U-shaped relationship between DA and performance emerges [145–150]. Rational inattention, as we have presented it, predicts that precision, and therefore performance, simply increase with DA. However, it is reasonable to assume that the encoding machinery has fundamental limits to its precision, so that arbitrary increases in DA may wildly increase estimated precision without an accompanying and equal increase in true precision. This will lead to precision miscalibration, and increases in DA over this range will now worsen the miscalibration and thus worsen performance. In Supporting Information, we derive this result analytically, and show that when true precision is bounded, the relationship between DA and performance takes the shape of an inverted U. This effect may also explain a number of apparent inconsistencies in the experimental literature. Notably, Beeler et al. [31] have found that chronically hyperdopaminergic mice were more willing to select high-cost levers in a two-armed bandit task than wild-type mice. Computational modeling of this experiment suggested that DA promotes exploration (rather than exploitation), which, as the authors note, may be explained by the U-shaped effect of DA. Similarly, behavior consistent with a *slower*, rather than faster, clock has been reported with optogenetic stimulation of midbrain DA neurons in mice [151] (but see [152, 153]), as well as in Parkinson’s patients who were off medication during training but on medication during testing on a separate day [40]. Whether these seemingly inconsistent findings are owed to the U-shaped effect of DA and its manipulation at non-physiological levels remains to be examined.

In restricting our analysis to the effects of tonic DA in reinforcement learning and interval timing, we have disregarded a wide array of experimental findings on DA. For instance, DA exhibits a distinct phenomenon of ‘ramping’ over the course of a single trial in a number of reinforcement learning tasks, such as goal-directed spatial navigation [154], bandit tasks [32], and timing of movement initiation [152], but not in others, such as classical conditioning tasks [1, 155–157]. These ramps, which occur on the timescale of seconds, are primarily observed during operant tasks, with a rapid return to baseline after task completion (e.g., during the post-reward delay). A natural question, then, is whether this differential in the average DA level before vs. after task completion mediates the effects of controllability on clock speed. Some authors have indeed interpreted these ramps as a ‘quasi-tonic’ signal [158], while others have argued in favor of an RPE interpretation of ramps, similar to that of phasic DA signals [153, 159–161]. DA has also been implicated in mediating spatiotemporal credit assignment in reinforcement learning tasks [162], and DA’s roles in working memory [163–166], spontaneous movement [167, 168], impulsivity [169–174], creativity [175, 176], and other domains have been the subject of great interest as well.

Second, seeking a computational theory of tonic DA necessarily introduced a number of simplifications. For instance, DA’s effects vary by receptor subtype: In the basal ganglia, which implements reinforcement learning models, the neurons primarily expressing either D1 or D2 receptors largely segregate anatomically into two separate pathways (the ‘direct’ and ‘indirect’ pathways, respectively, which later converge) [177] and seem to serve opposite purposes [178, 179]. DA bursts primarily potentiate D1 synaptic weights and depress D2 synaptic weights, and vice versa for DA dips [180]. Furthermore, the opposing effects of phasic DA on D1 and D2 receptors seems to extend to tonic DA as well [27, 181, 182]. On the other hand, based on pharmacological studies using D1 and D2 antagonists, the interval timing effects of DA seem to be primarily D2-mediated [113, 183], although recent work has highlighted a role for D1 as well [56, 113, 183–185]. While computational theories should transcend specific implementations [186], a computationally complete picture of DA will likely need to account for receptor heterogeneity [4, 27]. DA’s effects similarly vary by projection site (with broad projections across cortical and subcortical regions [187, 188]) and enzymatic activity [6, 189], and the predictions of the rational inattention framework cut across this diversity. For example, consider tonic DA’s role in the exploration-exploitation trade-off. Recent work has shown that DA’s action in prefrontal cortex modulates an exploration strategy referred to as ‘directed exploration,’ in which uncertainty about a reward source confers a value bonus, thus making the source more likely to be sampled. Striatal DA, on the other hand, has been linked to ‘random exploration,’ in which agents modify the stochasticity of their choices according to the total uncertainty in the environment [6, 190]. How these empirical findings may fit into a broader theory of tonic DA will be the subject of future work.

Third, in adopting the rational inattention framework, we have defined the ‘cognitive cost’ to be the cost of reducing the uncertainty in the world. This is far from the only cognitive cost an animal must pay in a task, and indeed different costs may outweigh others depending on the task. For a theory of tonic DA, our attentional cost function was informed by the empirical evidence linking DA to precision. However, a more complete optimization problem will need to incorporate capacity costs [e.g., 191], computation costs [192], interference costs [193], metabolic costs [194], and others [195]. How these different factors influence the optimization problem, and whether and how they interact with DA, remain to be examined.

Finally, this analysis does not preclude other factors from influencing the precision of encoding or the final estimate of decoding. Using reinforcement learning as an example, the volatility of an environment [196] and the stochasticity of reward sources [197] should affect the learning rate, but it is not clear that these quantities are reflected by the DA signal and rather are likely under direct cortical control [196, 198, 199] (but see [200, 201]). We have presented here the base case where, holding all else constant, average reward controls encoding and decoding in a very straightforward way. In more realistic environments where all else is not held constant, average reward will be one of a number of factors influencing encoding and subsequent decoding.

At first glance, the functions of DA seem to vary across processing stage and modality. We have shown how seemingly unrelated behaviors—such as modulation of the speed of an internal clock and learning from positive feedback—can be traced back to similar computations under the unifying principle of rational inattention.

## Methods

### Precision and gain modulation of value in reinforcement learning

In the Results, we argue that manipulating estimated precision has the apparent effect of controlling the gain of action values. Here, we describe this prediction concretely.

Standard models of action selection posit that the probability of selecting action *A_i_* with estimated value 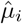 follows a softmax function [105, 106]:

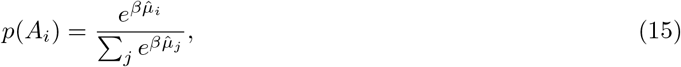

where *β* is referred to as the inverse temperature parameter. Recent influential models have argued that DA modulates the gain of the values 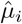, possibly by controlling the inverse temperature parameter *β* [27, 30, 202]. For simplicity, we examine the case of two actions, *A_l_* and *A_s_*, associated with large and small reward, respectively. Eq. (15) then reduces to

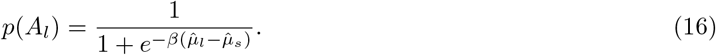

Importantly, the probability of selecting the large reward (exploitation) depends on the difference between the reward magnitudes and not on the absolute magnitudes themselves. As the quantity 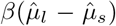 decreases, *p*(*A_l_*) decreases. Hence, any manipulation that results in a decrease in the estimated difference will encourage exploration over exploitation. Gain control is conventionally viewed as acting through the inverse temperature parameter *β*, which serves to amplify or reduce the influence of the difference 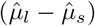 on the animal’s behavior. In this simple case, modifying *β* can be thought of as modifying choice stochasticity (see [203, 204] for more precise formulations). However, the same effect can be achieved by amplifying or reducing the estimated difference 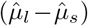 directly. Under Bayes’ rule, manipulating the likelihood precisions modulates the resulting difference in posterior means, thus implementing gain control on action values.

### Computational model of post-reward delay

We consider now models in which animals make their choices by simply maximizing the average reward over the entire task [205, 206]. In particular, Blanchard et al. [139] propose a discounting-free model of intertemporal choice, in which the animal maximizes non-discounted average reward (total reward divided by total trial duration). Thus the value *v_c_* of each choice *c* can be written as

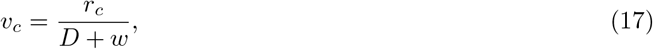

where *r_c_* is the actual reward for choice *c*, *D* is the pre-reward delay, and *w* is the estimated post-reward delay, which is a free parameter fit by behavior. Importantly, this model is mathematically translatable to hyperbolic discounting models, which also have a single parameter (in this case, the temporal discount factor *k*):

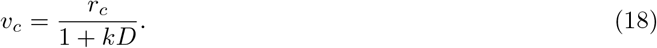

Thus, as the authors note, the data is fit by both models equally well, and cannot be used to arbitrate between them. (The authors argue against the discounting model elsewhere in their study, with Eq. (17) simply serving to examine the magnitude of *w* against the true post-reward delay.)

### Simulation details

#### Reinforcement learning

For the exploration-exploitation trade-off, we have chosen *κ* = 0.1, and reward magnitude of 1. Baseline DA was set to 0.9, and prior precision to 10. In all conditions, average reward was set to the DA level. Action selection was implemented using the softmax function in Eq. (15), with *β* = 4 (Fig. 3B). For learning from positive and negative feedback, we have chosen *κ* = 0.1, and reward magnitudes of 1 and 0 to reflect reward and punishment, respectively. To model the effect of average reward, reported by DA, on prior expectations in a new context, we set the prior mean to equal the DA level. (Note that, in general, the prior mean may be a combination of the average reward and recently experienced stimuli, weighted by their precisions.) Prior precision was arbitrarily set to 5. Accuracy was operationally defined as the area under the posterior distribution closer to the correct outcome (1 for rewards, 0 for punishments; i.e., the area to the right of and to the left of 0.5, respectively). Relative accuracy is the difference between these two areas (Fig. 3D).

#### Interval timing

For both experiments, our objective-subjective mapping is *m* = *η* log(*μ*+1) with subjective precision *l* = 1 to capture Weber’s law (see Supporting Information for a derivation). Average reward is equal to the DA level. For the central tendency, *κ* = 0.05*s*^−1^, and DA = 0.2 and 1 for low and high DA levels, respectively. By visual comparison with Shi et al. [41], we approximated the prior standard deviation to be 2.5 times smaller than the difference between presented durations. Prior precision is the inverse square of the prior standard deviation (Fig. 4B). For the speed of the internal clock, *κ* = 0.1*s*^−1^, DA = 0.8, 1, and 1.2 for the low, baseline, and high DA conditions, respectively (Fig. 4D).

#### Controllability

For both experiments, we set average reward to be equal to the DA level. Rate of timing-contingent rewards was set to 0.01, and *κ* = 0.1*s*^−1^. Note here that the reward incentive *R* formally refers to the subjective value of the reward, rather than its objective value. We have generally treated these quantities as interchangeable, as, under normal circumstances, they are monotonically related and therefore affect precision in the same qualitative way (Eq. (10)). However, one can specifically disentangle these two by manipulating satiety or baseline reward availability, as in Bizo and White [134]. Indeed, the incentive to accumulate rewards in satiated states is very low; the incentive to accumulate the same amount of reward but at near starvation is extremely high. To capture this distinction, we set *R* to be a decreasing function of baseline reward availability (more precisely, we set them to be inversely related). Note that this effect can also be captured by taking subjective reward to be a concave function of objective reward, following convention [207, 208]. This reflects the idea that the added benefit of an extra unit of reward decreases as more rewards are accumulated (Fig. 5B,C). For post-reward delays, we arbitrarily set average reward to be 1, with a baseline of 0.7 for the post-reward delay (Fig. 6C).

#### Experimental predictions

Small, medium, and large rewards had magnitude 5, 7.5, and 10, respectively. Precision under high and low controllability was set to 1 and 0.1, respectively. Action selection was implemented using the softmax function in Eq. (15), with *β* = 1 (Fig. 7). RPEs were computed as the difference between received reward (5 or 10) and the expected reward for the arm (7.5) (Fig. 7B). To simulate gain control, we set the inverse temperature parameter to *β* = 1 for low DA and *β* = 10 for high DA (Fig. 7C).

## Acknowledgements

The authors are grateful to Rahul Bhui for comments on the manuscript.

## Author contributions

J.G.M. and S.J.G. developed the model. J.G.M., L.L., and S.J.G. contributed to the writing of the paper. J.G.M. and L.L. made the figures. J.G.M. analyzed and simulated the model, and wrote the first draft.

## Data and code availability

Source code for all simulations can be found at www.github.com/jgmikhael/rationalinattention.

## Supporting Information

## 1 The relationship between reward incentive and precision

In the Results section, we derived the relationship between the reward incentive *R* and the optimal precision *λ*^*^ (Eq. (10) in the main text):

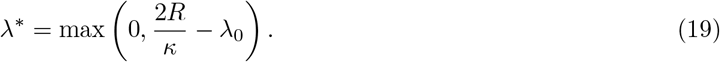

This relationship is piecewise linear (Fig. S1): When the incentive *R* is small, the precision *λ*^*^ is zero, and only after *R* becomes sufficiently large does *λ*^*^ begin to increase. Intuitively, the agent’s performance depends on its posterior precision, which is the sum of its prior and likelihood precisions. Under rational inattention, the agent seeks to ‘match’ its posterior precision with the reward incentives (divided by the information cost). When the prior precision alone achieves—or exceeds—the optimal posterior precision given the reward incentive, there is no need to attend to the task and collect new (costly) information, so *λ*^*^ is simply zero (horizontal segment of piecewise linear function). On the other hand, when the reward incentive calls for a higher posterior precision than the prior precision, the agent should make up the difference by attending to the task (increasing ray of piecewise linear function). The point at which the two pieces of the function intersect corresponds to the value of *λ*_0_ that exactly matches the reward incentive.

**Figure S1:**
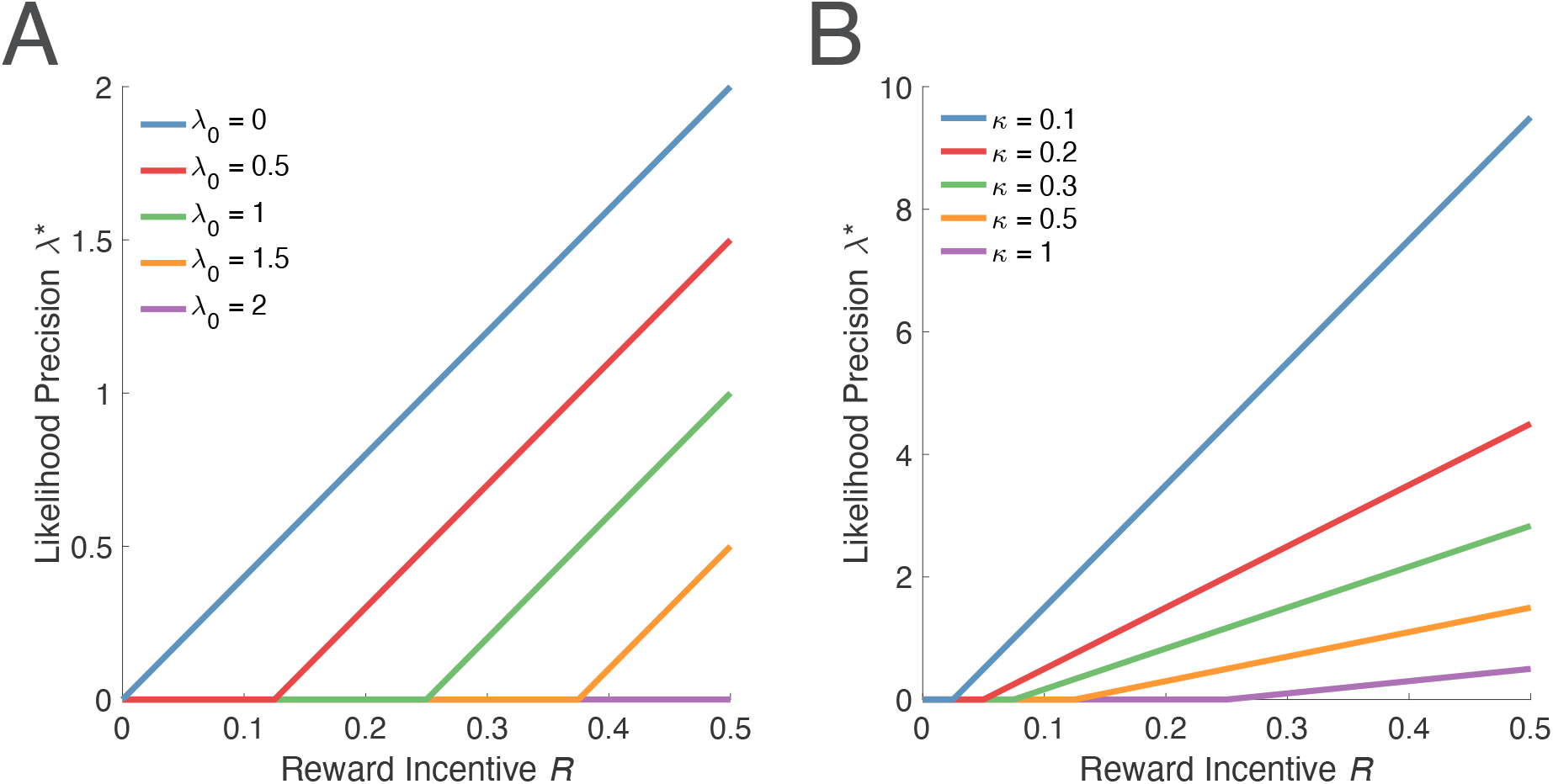
Relationship between reward incentive and likelihood precision under different levels of prior precision and information cost. The relationship between *λ*^*^ and *R* is piecewise linear: When *R* is small, the agent is not sufficiently incentivized to attend to the task and relies only on its prior knowledge. When *R* is sufficiently large, the agent linearly increases *λ*^*^ with *R*. (A) Increases in *λ*_0_ shift the piecewise linear function to the right. (B) Increases in *κ* shift the function to the right and decrease the slope. Simulation details: We have fixed *κ* = 0.5 and *λ*_0_ = 0.5 in (A) and (B), respectively.

It is straightforward to consider how changes in *λ*_0_ and *κ* affect the relationship between *R* and *λ*^*^. For larger values of *λ*_0_, the agent must be incentivized more before it begins to attend to the task (the point at which *λ*^*^ begins to increase shifts to the right). But after this point, a unit increase in *R* will increase *λ*^*^ by the same amount regardless of *λ*_0_ (same slope of increasing ray with different *λ*_0_; Fig. S1A). Similarly, for larger values of *κ*, larger reward incentives *R* will be needed for the agent to attend to the task. However, because precision depends on the ratio between *R* and *κ*, a unit increase in *R* will have a weaker effect on *λ*^*^ when *κ* is large (shallower slope; Fig. S1B).

## 2 The scalar property as a consequence of rational inattention

We outline here a derivation of the scalar property (Weber’s law) based on rational inattention principles. The purpose of this derivation is to provide a mathematically well-specified model of timing noise (or likelihood precision). Quantitatively, this derivation will control the observed increase in timing noise for larger intervals, but our main effects (central tendency and changes in clock speed) do not depend on this derivation, and only depend on the assumption that the control of precision be implemented via changes in clock speed.

In Eq. (10), we derived the relationship between reward, information cost, and precision. When the temporal task is not highly conditioned, *λ*_0_ C 0, and we can simply write

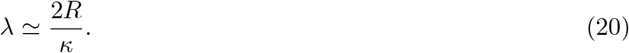

(In highly conditioned tasks, on the other hand, *λ*_0_ becomes large, the central tendency effect becomes prominent, and the scalar property is not guaranteed to apply [42].)

In interval timing tasks, rewards are subject to hyperbolic discounting [209–211]:

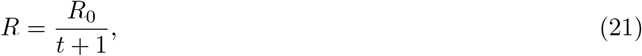

where *R*_0_ is the undiscounted reward, which in our framework is reported by DA, and *t* is time. Following previous work [212], we assume that the cost of collecting information (reducing noise) scales with the attended duration, and we take this scaling to be linear:

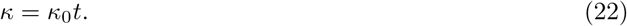

Here, *κ*_0_ represents the attentional cost per unit time. It follows then that

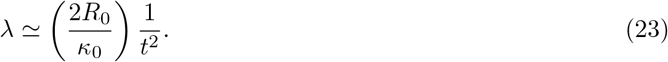

This is the scalar property, with Weber fraction given by:

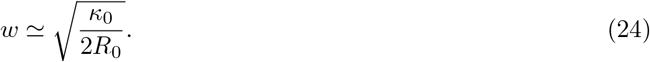

This leaves open the question of how the scalar property is actually implemented. One approach is to take a concave (e.g., logarithmic) mapping of objective time to subjective (psychological) time:

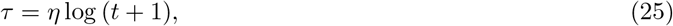

where *τ* represents subjective time, and *η* is a scaling factor, which can be interpreted as the pacemaker rate of the internal clock [213–215]. We hold subjective precision *l* constant. Bayesian update occurs in subjective time, and is subsequently mapped back to objective time during decoding.

For any desired precision *λ*, we can then find *η* to achieve this precision. In particular, by deriving the relationship between subjective and objective precision:

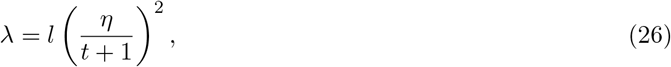

it follows that

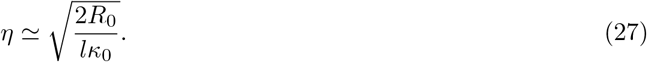

Hence, average reward *R*_0_ controls precision *λ* by setting *η* (e.g., see Fig. S2). In particular, the speed of the clock increases with reward (*R*_0_) and decreases with cost (*κ*_0_).

Interestingly, this framework predicts that the scalar property should not apply over all time ranges, but rather, the ratio 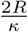 eventually becomes small enough that *λ*_0_ cannot be disregarded, and eventually, 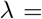 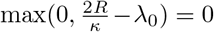. After that point, it is no longer worth using this mechanism to time. Instead, a different timing mechanism in which *κ* does not increase linearly should apply. This may explain why interval timing only applies over the seconds-to-hours range [48].

A second prediction of this derivation is that changes in precision will be more evident under changes to the reward rate, rather than the reward magnitude. To see this, assume the timed duration at baseline is *T*, and the average reward is increased from *R* to *nR* > *R*. If this change occurred by increasing reward *rate*, then the duration to be attended to before each reward is delivered becomes 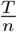, and the magnitude of each reward remains *R*. Hence for each reward, both the cost and reward discounting scale by 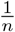. On the other hand, if this change occurred by increasing reward *magnitude*, then the magnitude increases to *nR*, but the duration attended to remains *T*. Hence cost and reward discounting are unchanged. Let us compute the new precisions, *λ_r_* and *λ_m_*, respectively:

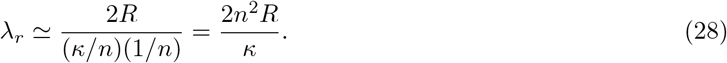

On the other hand,

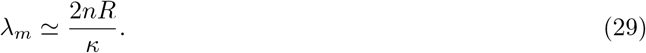

Therefore, *λ_r_* > *λ_m_*.

In fact, subjective reward magnitudes are concave with objective magnitudes, rather than linear [207, 208].

This further amplifies the difference between the two precisions: In this case,

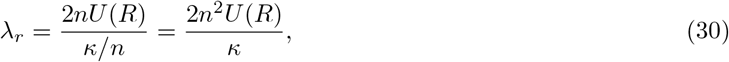

where *U* (*R*) represents the subjective value of reward *R* (the utility of *R*), which is a concave function. On the other hand,

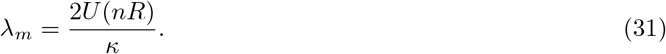

This result predicts that *λ_r_* ≫ *λ_m_*. Indeed, changes in reward rate have much more profound effects on interval timing than changes in reward magnitude [e.g., 117, 122].

## 3 Time cell rescaling

In the previous section, we showed that average reward *R*_0_, which in our model is reported by DA, controls precision *λ* by setting the pacemaker rate *η*. As shown in Fig. S2, the temporal receptive field surrounding any given objective time tightens under higher DA levels.

**Figure S2:**
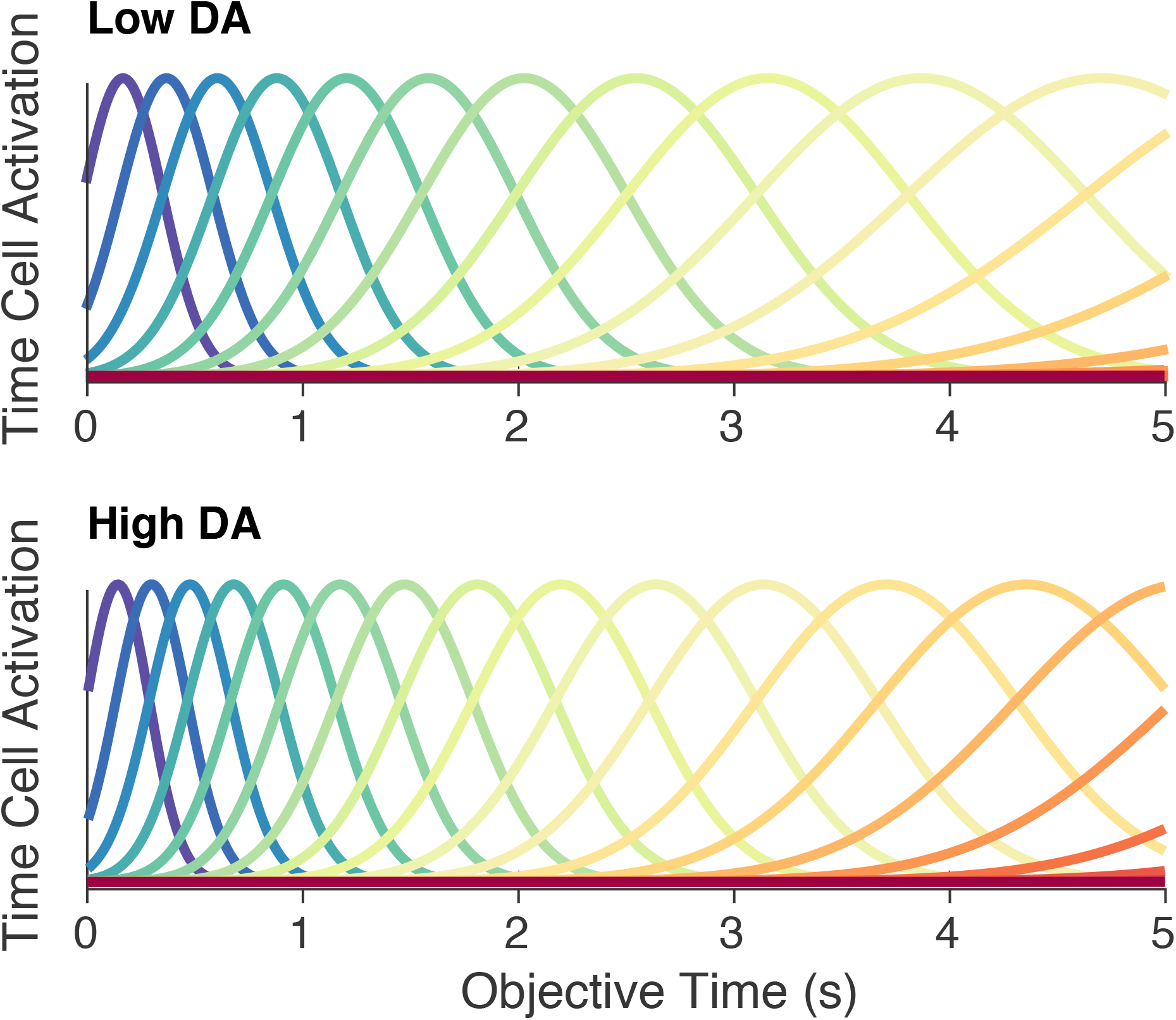
Time cell receptive fields rescale to increase precision. Mathematically, manipulations of precision correspond to changing the mapping between subjective and objective time. Simulation details: We have chosen *κ*_0_ = 0.05*s*^−1^, *λ* = 1 in subjective time, and DA = 1 and 1.5 for the low and high DA conditions, respectively.

## 4 Interval timing in discrimination tasks

Interval timing is typically studied using production tasks [125], such as the experiments considered in the main text, or discrimination tasks [216]. In discrimination tasks, subjects (or animals) learn to respond differently to short-duration stimuli and long-duration stimuli. Intermediate-duration stimuli are then presented during probe trials, and for each, the subjects must respond with the short-duration response or the long-duration response. Subsequent changes in their response profiles following certain manipulations (e.g., administration of drugs) allow us to infer changes in their internal timing system.

**Figure S3:**
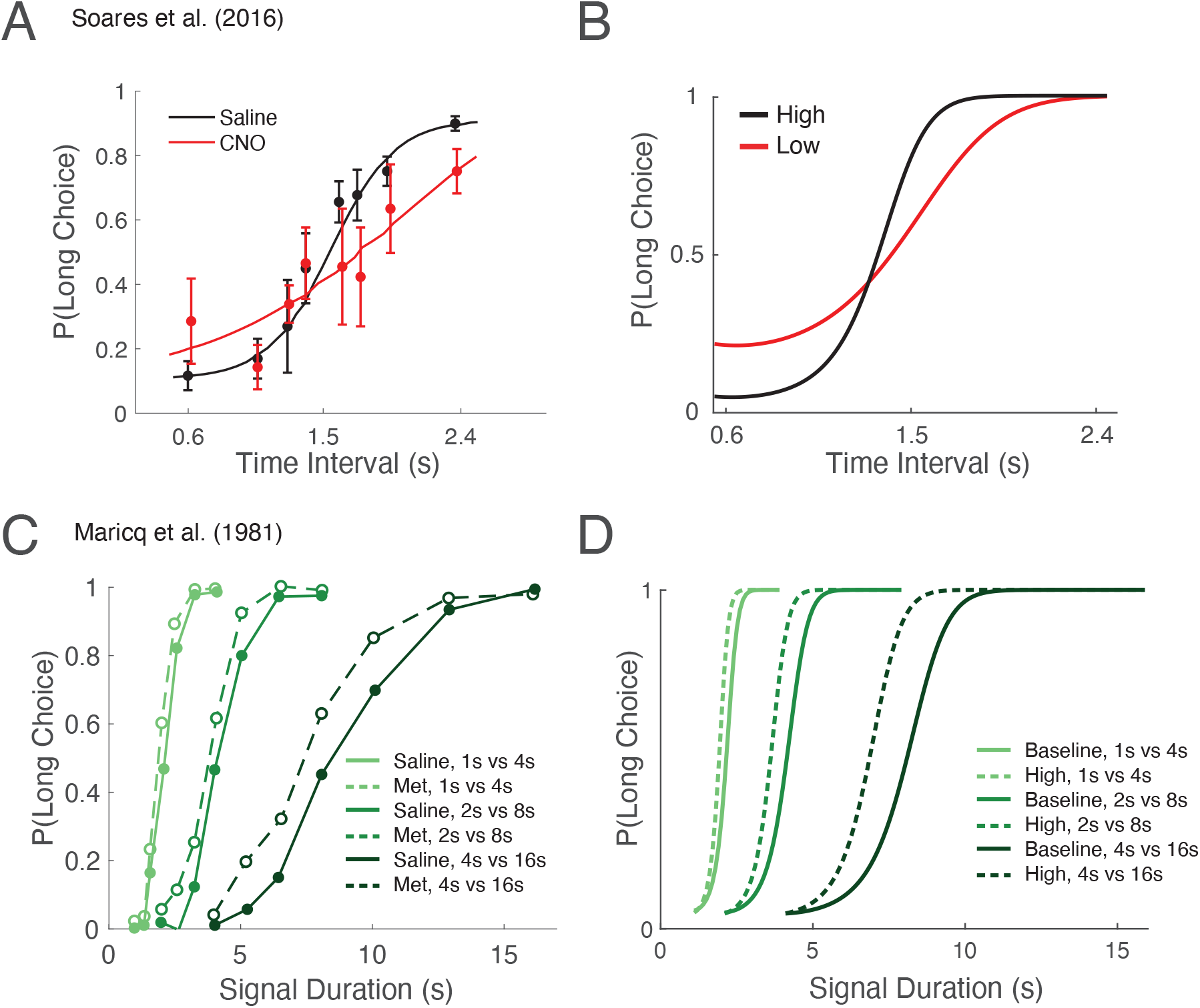
Rational inattention and interval timing in discrimination tasks. (A) Soares et al. [18] trained mice on a temporal discrimination task in which they reported intervals of variable duration as either shorter or longer than 1.5 seconds. When DA activity was pharmacogenetically suppressed, the discrimination curve flattened. p(Long Choice): probability of judging a duration as long; CNO: clozapine N-oxide. Figure adapted from Soares et al. [18]. (B) Our model recapitulates this effect: Under rational inattention, high DA increases precision, which mitigates Bayesian migration and improves discrimination. Note here, however, that motivation is a confounding factor. (C) Maricq et al. [15] have shown that, in rats, acute administration of methamphetamine (DA agonist) during testing in discrimination tasks with various time pairs biased estimation toward the ‘long’ response. Pairs of solid and dashed curves represent conditions where the short and long durations were 1 and 4 seconds (light green), 2 and 8 seconds (green), and 4 and 16 seconds (dark green). This discrimination paradigm controls for motivation, a potential confound in reproduction tasks (Fig. 4C). Figure adapted from Maricq et al. [15]. (D) Our model recapitulates this effect: High DA at decoding increases the speed of the clock, which biases estimation of duration toward longer responses. Simulation details: We have chosen *κ*_0_ = 0.2*s*^−1^, and DA levels of 0.3, 0.7 and 0.8 for low, baseline, and high conditions, respectively. Average reward was set to the DA level.

## 5 Interval timing under average reward manipulations

**Figure S4:**
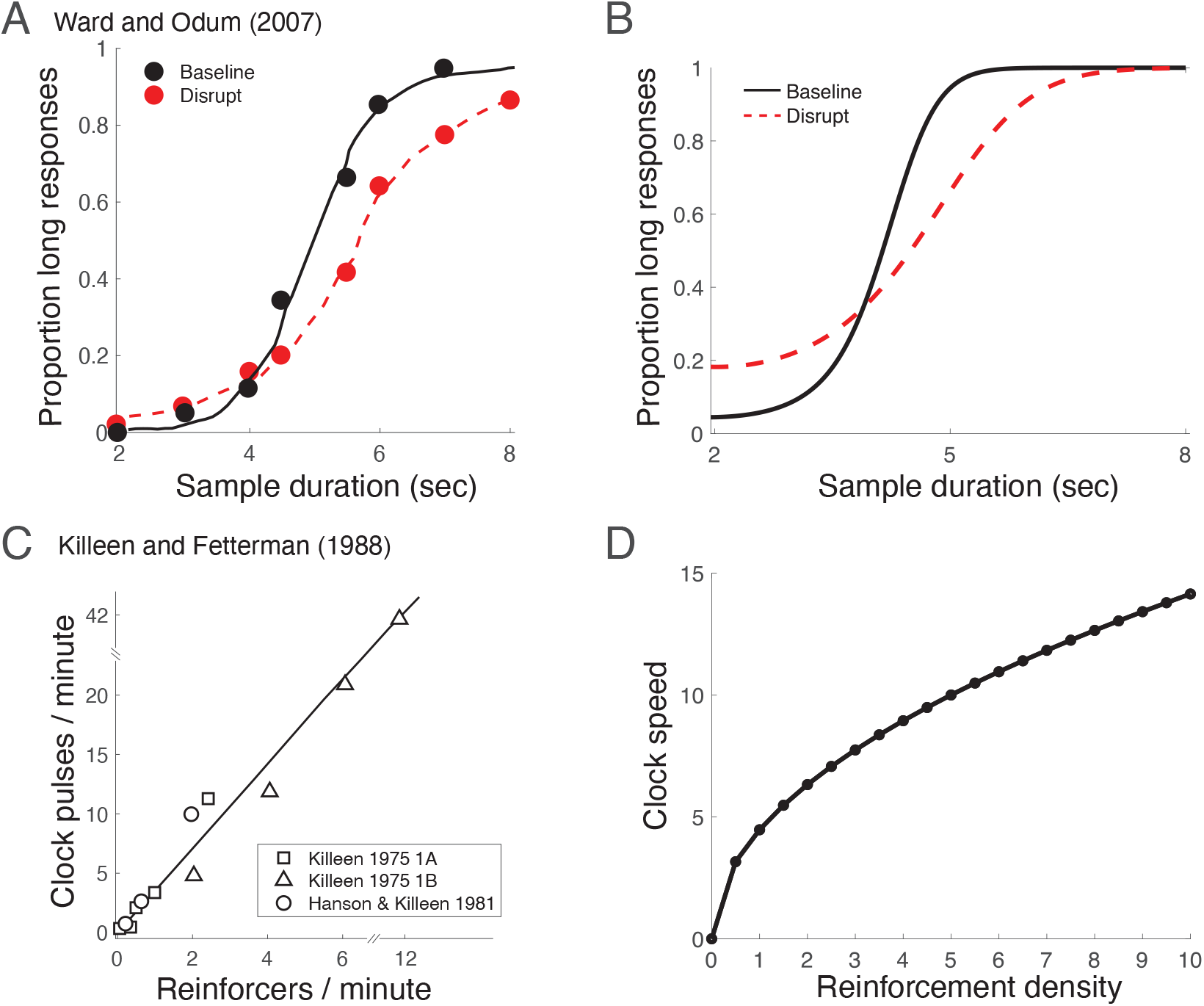
Effects of average reward and satiety on interval timing. (A) Ward and Odum [110] trained pigeons on a temporal discrimination task in which they reported intervals of variable duration as either shorter or longer than 5 seconds. When the animals were prefed, the discrimination curve flattened. Figure adapted from Ward and Odum [110]. (B) Our model recapitulates this effect: Under rational inattention, high reward incentives increase precision, which mitigate Bayesian migration and improve discrimination. Note here, however, that motivation is a confounding factor. (C) Killeen and Fetterman [117] analyzed data from three experiments in which reinforcer rates were varied, and timing behavior was measured. The authors fit behavior to a model in which the speed of the internal clock could be manipulated. They showed that the clock speed increased as the reinforcer rate increased. Clock pulses/minute: Model parameter representing clock speed. Figure adapted from Killeen and Fetterman [117]. (D) Our model recapitulates this effect: Under rational inattention, high average reward increases precision. This results in a faster internal clock. Simulation details: We have chosen *κ*_0_ = 0.2*s*^−1^, and DA levels of 0.3 and 0.7 for high and low satiety, respectively. Average reward was set to the DA level.

## 6 Inverse U-shaped relationship between DA and performance

We assume here that true precision *λ* cannot increase arbitrarily, and instead is bounded by *λ* ≤ *L*, for some constant *L*. We will show that if the estimated precision *λ′* can increase beyond *L*, then performance will follow an inverse U-shaped curve with DA level. Define *d* as the desired (or targeted) precision, controlled by DA. Then *λ* = min(*d, L*), and *λ′* = *d*.

Combining Eqs. (13) and (14), and knowing that *c* = *λ′/λ*, we can rewrite the marginal error as a function of *d*, which is monotonic with DA:

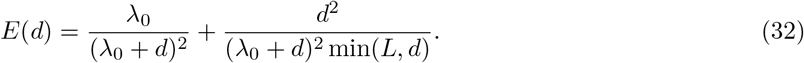

When *d* ≤ *L*, 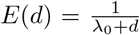 Hence, over this domain, error will decrease as DA increases. However, when DA increases enough for *d* to surpass *L*,

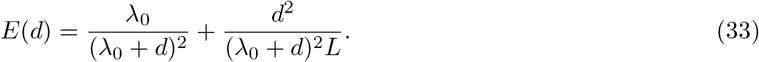

Taking the partial derivative of *E* with respect to *d*,

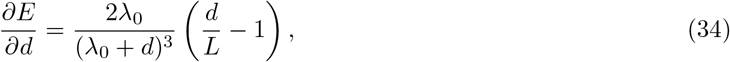

which is positive when *d* > *L*. Hence, error decreases with DA until true precision reaches the limit *L*. After this point, increases in DA increase error (Fig. S5). In other words, the relationship between DA and performance takes the shape of an inverted U.

**Figure S5:**
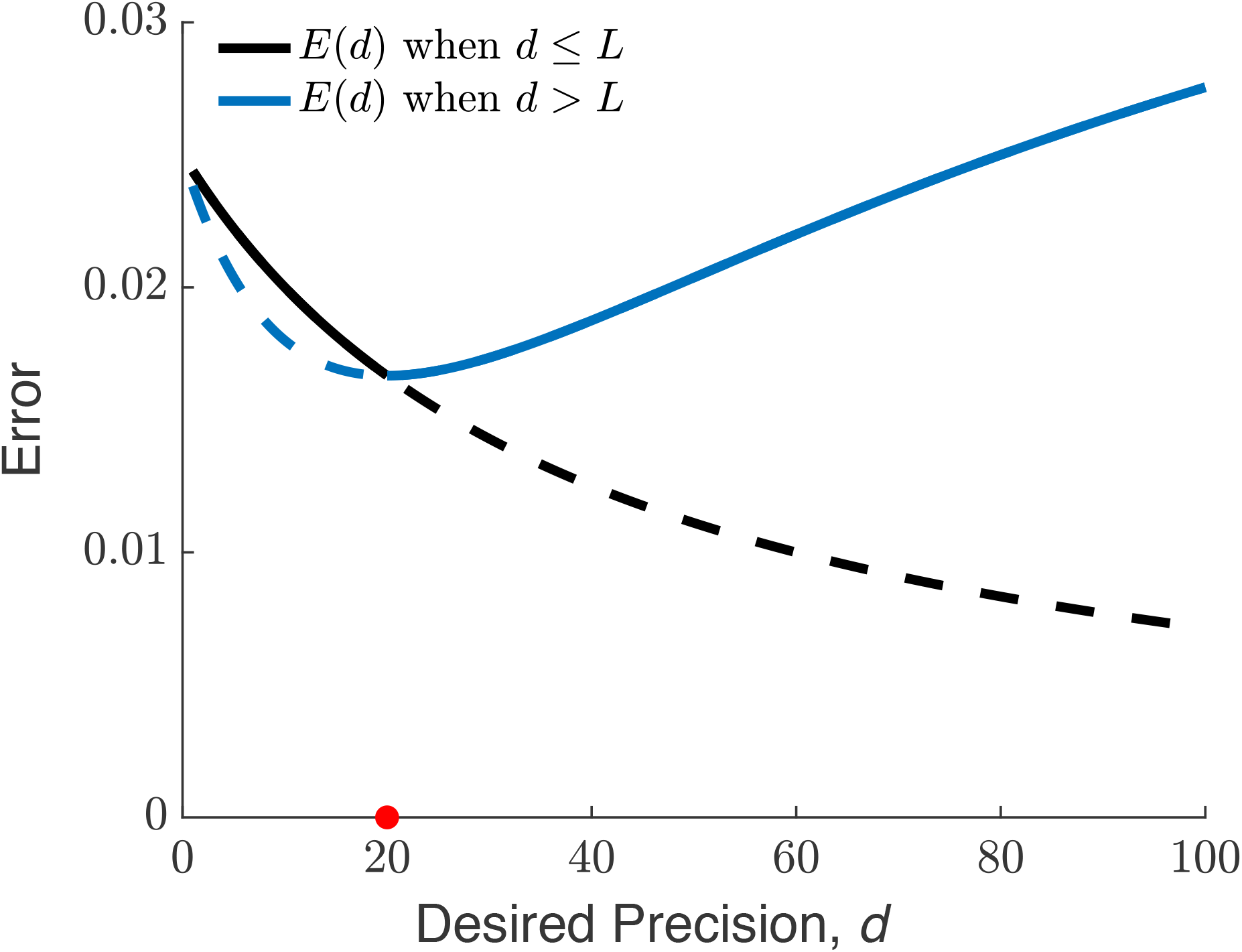
When true precision is bounded from above, the relationship between DA and error is U-shaped. When DA is low, increases to DA will improve both true and estimated precision, which improves performance. However, when DA increases beyond the capacity of true precision to increase, precision miscalibration will cause performance to worsen (see ‘Precision miscalibration’ section). Because performance depends inversely on error, it follows that the relationship between DA and precision is inverse U-shaped. Dashed curves denote the error if precision were not bounded from above (black) and if true precision were fixed at *L* independent of the desired precision (blue). For illustration, we have chosen *L* = 20 and *λ*_0_ = 40. *d*: desired precision set by DA; *E*(*d*): error as a function of desired precision; *L*: limit to true precision, indicated by red dot.

